# Prostaglandins regulate invasive, collective border cell migration

**DOI:** 10.1101/821686

**Authors:** Emily F. Fox, Maureen C. Lamb, Samuel Q. Mellentine, Tina L. Tootle

## Abstract

While prostaglandins (PGs), short-range lipid signals, regulate cell migration, their mechanisms of action are poorly understood in collective migration. To address this, we use *Drosophila* border cell migration during Stage 9 of oogenesis. The border cells delaminate from the epithelium, and migrate collectively and invasively between the nurse cells. Pxt is the *Drosophila* cyclooxygenase-like enzyme responsible for all PG synthesis. Loss of Pxt results in both a significant delay in border cell migration during Stage 9 and an increase in cluster length compared to wild-type controls. Contributing to these phenotypes is altered integrin localization. Integrins are enriched on the border cell membranes, and this enrichment is lost in *pxt* mutants. Active integrins require interaction with the actin cytoskeleton. As we previously found PGs regulate the actin bundler Fascin and Fascin is required for border cell migration, we hypothesized PGs regulate Fascin to control integrins. Supporting this, loss of Fascin results in a *pxt*-like integrin localization, and dominant genetic interaction studies reveal that co-reduction of Pxt and Fascin results in delayed and elongated border cell clusters. Together these data lead to the model that PG signaling controls Fascin, and thereby integrins, to mediate on-time border cell migration and maintain cluster cohesion.

## Introduction

During invasive, collective cell migration multiple cells migrate as a group between other cells and/or through tissues. Such migration requires the coordination of numerous factors regulating cell migration including cell polarity, cytoskeletal remodeling, and notably, adhesion dynamics. Migratory cells must properly adhere and release from their substrate to promote migration (Grashoff *et al*., 2010; De Pascalis and Etienne-Manneville, 2017). Collective migration also requires maintaining communication and adhesions between all the migratory cells of the cluster (Mayor and Etienne-Manneville, 2016). While many factors regulating cell migration have been uncovered by studying single cell migration, *in vivo* most cell migration occurs as collective cell migration, including during embryonic development, regeneration, and cancer metastasis (Friedl and Gilmour, 2009; Scarpa and Mayor, 2016). Thus, it is critical to define the factors regulating collective cell migration.

One regulator of cell migration is prostaglandins (PGs) (Menter and Dubois, 2012). PGs are short-range lipid signals produced at their sites of action (Tootle, 2013). Cyclooxygenase (COX) enzymes convert arachidonic acid into the PG precursor, PGH_2_. This precursor is then acted upon by PG-type specific synthases to produce the different bioactive PGs. Each PG is secreted to bind and activate one or more G protein-coupled receptors to elicit different downstream signaling cascades to mediate various cellular and physiological outcomes. One PG, PGE_2_, is widely implicated in promoting cell migration both during development and cancer progression. Inhibition of COX activity or loss of PGE_2_ signaling alters cellular adhesion dynamics and blocks gastrulation in zebrafish (Cha *et al*., 2005; Cha *et al*., 2006; Speirs *et al*., 2010). Additionally, PGE_2_ regulates vascular maturation and angiogenesis (Ugwuagbo *et al*., 2019), homing of hematopoietic stem cells to their niche (North *et al*., 2007; Hoggatt *et al*., 2009), and macrophage migration (Digiacomo *et al*., 2015). Notably, the majority of these PG-dependent developmental migrations occur as collectives or groups of cells. PGs are also widely implicated in promoting cancer migration and metastasis (Menter and Dubois, 2012), both by functioning within the tumor cells and within the microenvironment (Li *et al*., 2015; Kobayashi *et al*., 2018). Cancer cells can migrate as both single cells and collectives (Friedl and Mayor, 2017; Pandya *et al*., 2017). Recent evidence suggests that collectively migrating cancer cells are more likely to establish metastatic tumors, are resistant to chemotherapies, and correlate with a poor prognosis (Giampieri *et al*., 2009; Alexander and Friedl, 2012; Khalil *et al*., 2017; Stuelten *et al*., 2018). As collective cell migration is important for normal development and contributes to cancer progression, and PGs regulate migration in both of these contexts, it is essential to establish a robust system for defining the mechanisms by which PGs regulate invasive, collective cell migration.

*Drosophila* oogenesis is an ideal *in vivo* model to uncover the roles of PGs (Tootle and Spradling, 2008; Spracklen and Tootle, 2015). Each female fly has two ovaries composed of chains of sequentially developing follicles or eggs (Spradling, 1993). Follicle development is divided into fourteen morphological stages. Each follicle is made up of 16 germline-derived cells, one posterior oocyte and 15 nurse cells. These germ cells are surrounded by a layer of somatic epithelial cells termed follicle cells. The follicle cells can be divided into different sub-types, including the outer follicle cells, stretch follicle cells, centripetal cells, and the border cell cluster, which includes both polar and border cells. *Drosophila* possess a single COX-like enzyme, Pxt (Tootle and Spradling, 2008). Loss of Pxt results in multiple defects during egg or follicle development and results in female sterility (Tootle and Spradling, 2008; Tootle *et al*., 2011; Groen *et al*., 2012; Spracklen *et al*., 2014; Spracklen and Tootle, 2015).

During *Drosophila* oogenesis, the process of border cell migration has been widely used to uncover conserved mechanisms regulating invasive collective cell migration (Montell, 2003; Montell *et al*., 2012). During Stage 9 (S9), 8-10 follicle cells differentiate into border cells, delaminate from the follicular epithelium, and collectively migrate from the anterior tip of the follicle, between the much larger nurse cells, to the nurse cell-oocyte boundary by S10A. During this migration, the follicle grows in size, the nurse cells become covered in squamous stretch follicle cells, and the outer follicle cells ultimately cover only the oocyte. The border cell cluster then migrates to the dorsal side of the oocyte and ultimately forms the micropyle, the structure through which the sperm enters to fertilize the egg (Montell *et al*., 1992). Thus, border cell migration is required for fertility.

Here we utilize border cell migration to uncover the roles of PGs in invasive, collective cell migration. Examination of S10 follicles reveals that PGs are required for regulating cluster morphology, as loss of Pxt results in elongated clusters with cells being left along the migration path. To uncover the cause of these defects, border cell migration was examined during S9. Loss of Pxt results in delayed border cell migration during S9 and aberrant, elongated border cell clusters. Knockdown of Pxt in the somatic cells results in delayed border cell migration, but the clusters are more compact. These findings suggest that PGs act in both the germline and the somatic cells to regulate border cell migration. We find one means by which PGs affect border cell migration is by regulating integrins. Loss of Pxt results in a decrease in membrane enrichment of integrins. A similar phenotype is observed when Fascin (*Drosophila* Singed), an actin bundling protein, is lost. As we previously found that PGs regulate Fascin to control actin remodeling within the germline (Groen *et al*., 2012; Spracklen *et al*., 2019) and Fascin is required for on-time border cell migration during S9 (Lamb *et al*., 2019), we postulated that PGs regulate Fascin to control border cell migration. Indeed, dominant genetic interaction studies reveal that co-reduction of Pxt and Fascin phenocopies loss of Pxt, resulting in delayed and elongated border cell clusters. Together these data lead to the model that Pxt produces PGs, which activate a signaling cascade to control Fascin, and thereby integrins, to mediate on-time border cell migration and maintain cluster cohesion.

## Results

### Pxt regulates border cell cluster morphology

A common means of assessing border cell migration is to determine if the cluster reaches the nurse cell-oocyte boundary by S10A. Prior work defining the roles of Pxt during *Drosophila* oogenesis reported that while the border cell cluster did reach the oocyte by S10A, the cluster had a long trail of cells remaining along the migration path (Tootle and Spradling, 2008). Extending from these studies, we sought to quantify the border cell defects when Pxt is lost.

For our analyses, we make use of two insertional *pxt* alleles – *EY03052* (*EY*) and *f01000* (*f*). Prior work characterizing these alleles has shown that *pxt^EY/EY^* exhibits a low level of *pxt* expression by both qRT-PCR and in situ hybridization (Tootle and Spradling, 2008). These same analyses revealed *pxt^f/f^* exhibited little to no *pxt* expression and immunoblotting revealed no protein product, suggesting *pxt^f^* is a null allele (Spracklen *et al*., 2014). Using these alleles, we assessed border cell migration at S10 by labeling the follicle cell nuclei, including the border cells, by immunofluorescence for Eyes absent (Eya). While the border cell clusters in *pxt* mutant follicles reach the nurse cell-oocyte boundary, the clusters are abnormal. Small groups or long chains of border cells are often left behind between the nurse cells in *pxt* mutant follicles (Figure 1B-C’ compared to A-A’, yellow arrows and bracket). Only 8% of wild-type follicles exhibit multiple border cells being left behind, while 76% of *pxt^f/f^*, 16% of *pxt^EY/EY^* and 48% of *pxt^EY/f^* mutant follicles have trailing border cells (Figure 1B-C’ compared to A-A’, and Supplemental Figure 1A). Specifically, we find that in *pxt^f/f^* follicles there is an average of 2.24 and in *pxt^EY/f^* follicles there is an average of 0.76 trailing border cells, compared to 0.11 in wild-type (Supplementary Figure 1A, p<0.0001). In addition, the number of border cells is increased when Pxt is completely lost; wild-type follicles exhibit an average of 5.2 border cells while *pxt^f/f^* follicles exhibit an average of 9.1 border cells (Supplementary Figure 1B, p<0.0001). Notably, the number of polar cells within the border cell cluster is not changed (data not shown). Together these data indicate Pxt contributes to border cell migration by regulating the border cell number and cluster cohesion.

**Figure 1:**
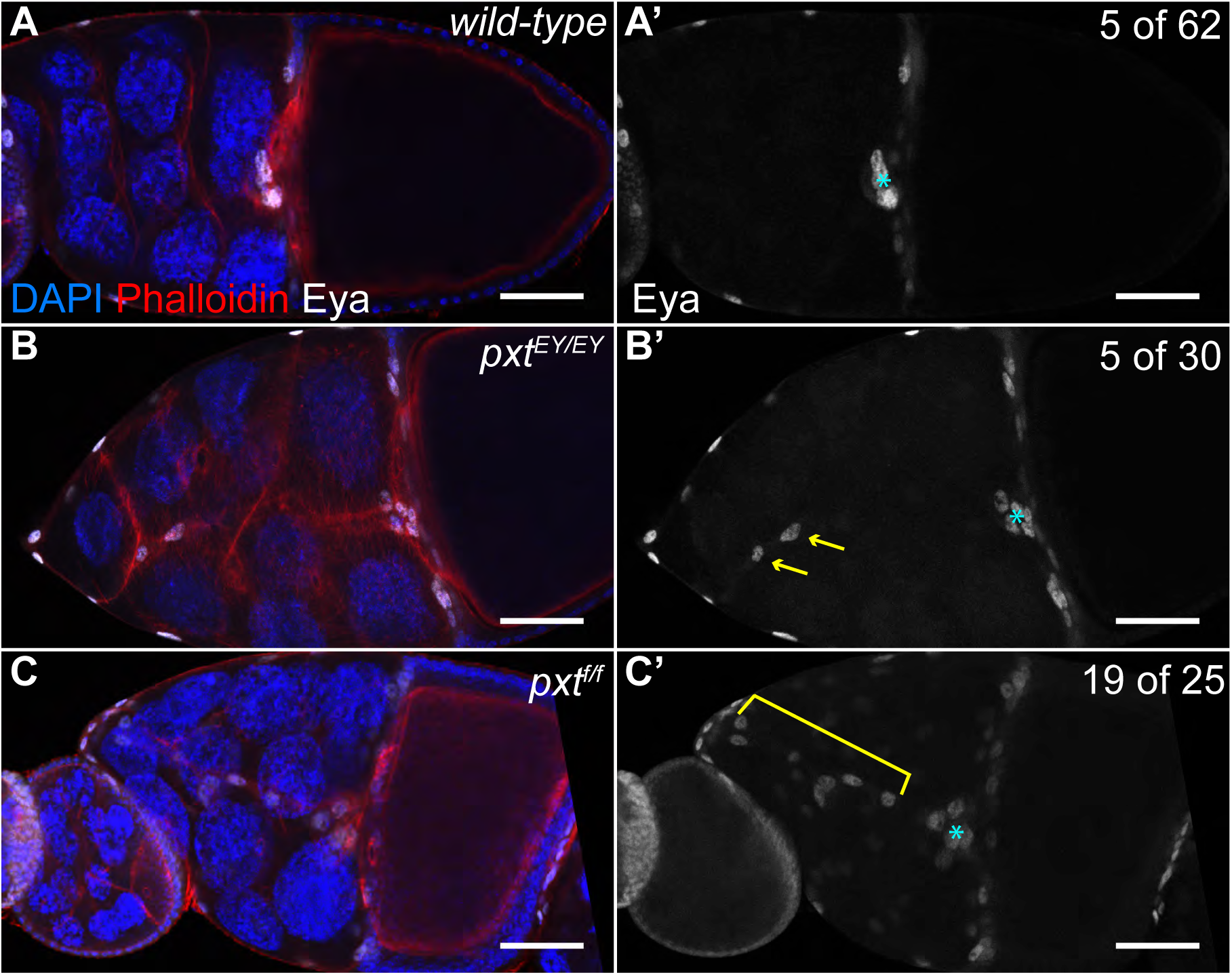
Prostaglandins regulate border cell cluster integrity. A-C’. Maximum projections of 3 confocal slices of S10 follicles of the indicated genotypes; anterior is to the left. A-A’. *wild-type* (*yw*). B-B’. *pxt^EY03052/EY03052^* (*pxt^EY/EY^*). C-C’. *pxt^f01000/f01000^* (*pxt^f/f^*). A-C. Merged images: Eyes absent (Eya), white; Phalloidin (F-actin), red; and DAPI (DNA), blue. A’-C’. Eya, white. The nuclei of the border, stretch follicle, and centripetal cells are marked by Eya staining. By S10, the intact border cell cluster is normally located at the nurse cell/oocyte boundary (A-A’, cyan asterisk). In *pxt* mutants, despite the majority of the cluster reaching the boundary (cyan asterisk), cells are often left behind (B-C’); the frequency of Stage 10 follicles exhibiting border cells left behind is indicated in the top right of panels A’-C’. These cells can exist as single cells or pairs of cells being left behind (B-B’, yellow arrows), or long continuous chains of cells being left behind (C-C’, yellow bracket). Scale bars = 50μm.

### Pxt is required for on-time border cell migration and maintenance of cluster morphology

To determine how the border cells defects arises when Pxt is lost, we examined border cell migration during S9. While live imaging is an ideal approach to define defects during the invasive, collective migration of the border cells (Prasad and Montell, 2007), *pxt* mutant follicles proved difficult to keep alive during the live imaging process (data not shown). Therefore, we assessed border cell migration from fixed immunofluorescence images. In wild-type follicles, the outer follicle cells (orange dashed line) are in line with the border cell cluster (green) throughout the migration (Figure 2A-B). Similarly, heterozygotes for mutations in *pxt* (*pxt-/+*) also exhibit on-time border cell migration (Figure 2C-D). Surprisingly, when either *pxt* allele is over the *MKRS* balancer chromosome, border cell migration is delayed (data not shown and Supplemental Figure 2); we speculate this is due to a genetic interaction with a mutation on the balancer chromosome and *pxt*. Loss of Pxt by either homozygosity for either mutant allele (data not shown) or transheterozygosity for both alleles (*pxt^EY/f^*) results in delayed border cell migration, as the border cells remain anterior to the outer follicle cells (Figure 2E-F).

**Figure 2:**
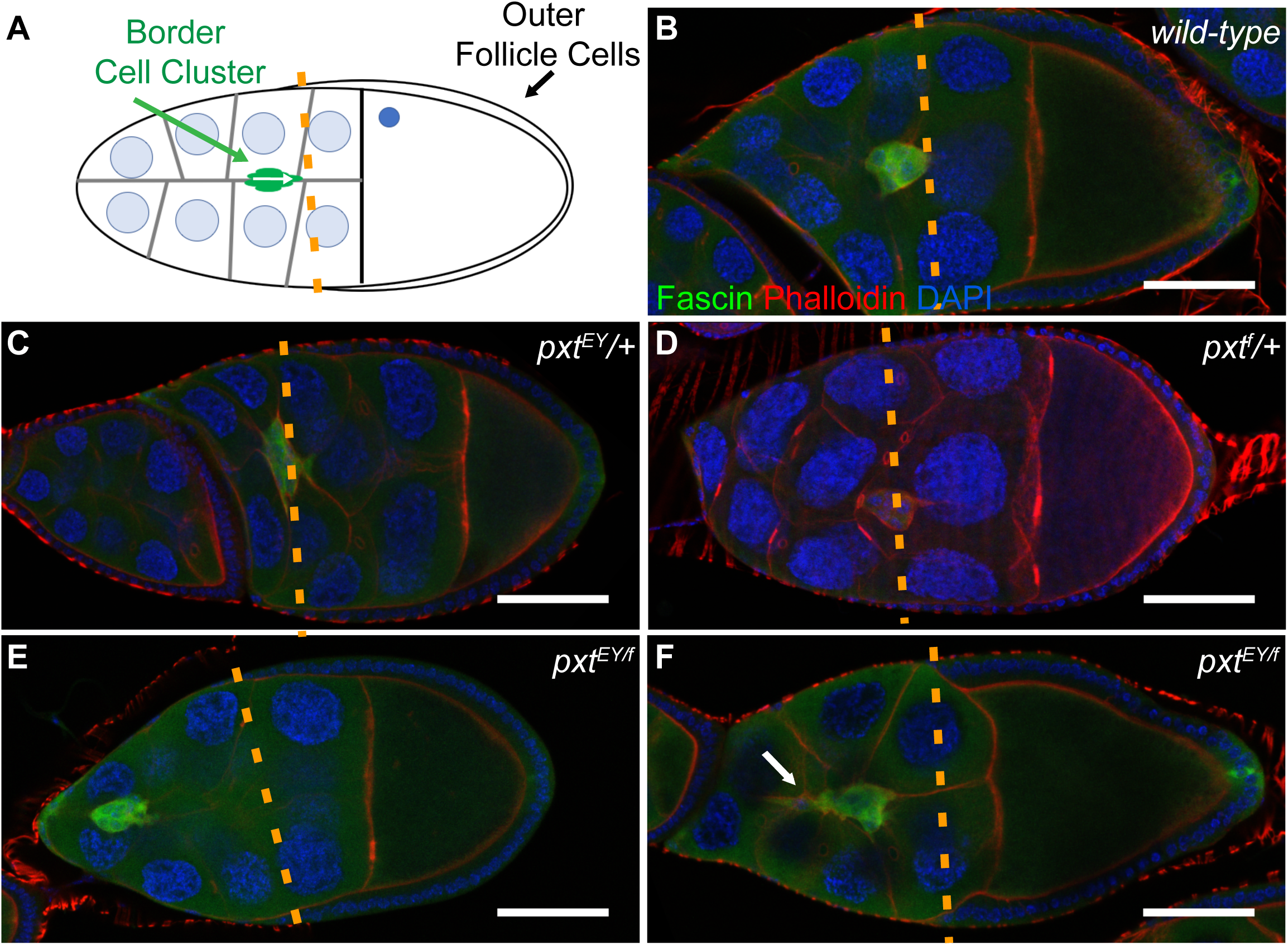
Prostaglandins are essential for on-time border cell migration during S9. A. Diagram depicting the wild-type alignment of the position of the border cell cluster (green; white arrow indicates direction of migration) with that of the outer follicle cells (orange dashed line) during S9 of oogenesis. B-F. Maximum projections of 3 confocal slices of S9 follicles of the indicated genotypes; anterior is to the left. B. *wild-type* (*yw*). C. *pxt^EY^/+*. D. *pxt^f^/+*. E-F. *pxt^EY/f^*. Merged images: Fascin, green; Phalloidin (F-actin), red; and DAPI (DNA), blue. Orange dashed lines indicate the position of the outer follicle cells. While the position of the border cell cluster is typically in line with the outer follicle cells in wild-type and heterozygous *pxt* mutant follicles (B-D, orange dashed line), when Pxt is lost the clusters are anterior to the outer follicle cells (E-F, orange dashedline). Additionally, the trans-allelic combination of the two *pxt* alleles often results in an elongated border cell cluster tail (F, white arrow). Scale bars = 50μm.

To further characterize the border cell migration defects during S9 we developed a quantitative method of assessing migration from fixed immunofluorescence images (Figure 3A). Specifically, we measure the distance the border cells have migrated from the anterior end of the follicle and subtract the distance the outer follicle cells are from the anterior of the follicle. We term this the migration index (units = μm). A migration index value of ∼0 indicates on-time migration, while negative values indicate delayed and positive values indicate accelerated migration. On average wild-type clusters exhibit a migration index of 2.93μm with a normal distribution between ∼25 and −25μm (Figure 3B). Loss of Pxt results in a significant delay in border cell migration, between ∼0 and −50μm, and exhibits an average migration index of - 23.12μm for *pxt^EY/EY^* and −28.26μm for *pxt^f/f^* follicles (Figure 3B, p<0.0001). Additionally, transheterozygotes of the two alleles (*pxt^EY/f^*) exhibit a migration index of −19.87μm (Figure 3B, p<0.0001). The negative migration indices could result from either delayed border cell migration or altered outer follicle cell distance. To distinguish between these possibilities, we plotted the follicle cell length versus the distance of the outer follicle cells for wild-type (red) and *pxt^EY/f^* (green), and find that they exhibit a similar slope, indicating outer follicle cell behavior is normal in *pxt* mutants (Supplemental Figure 3). Together these findings indicate that Pxt is essential for on-time border cell migration during S9.

**Figure 3:**
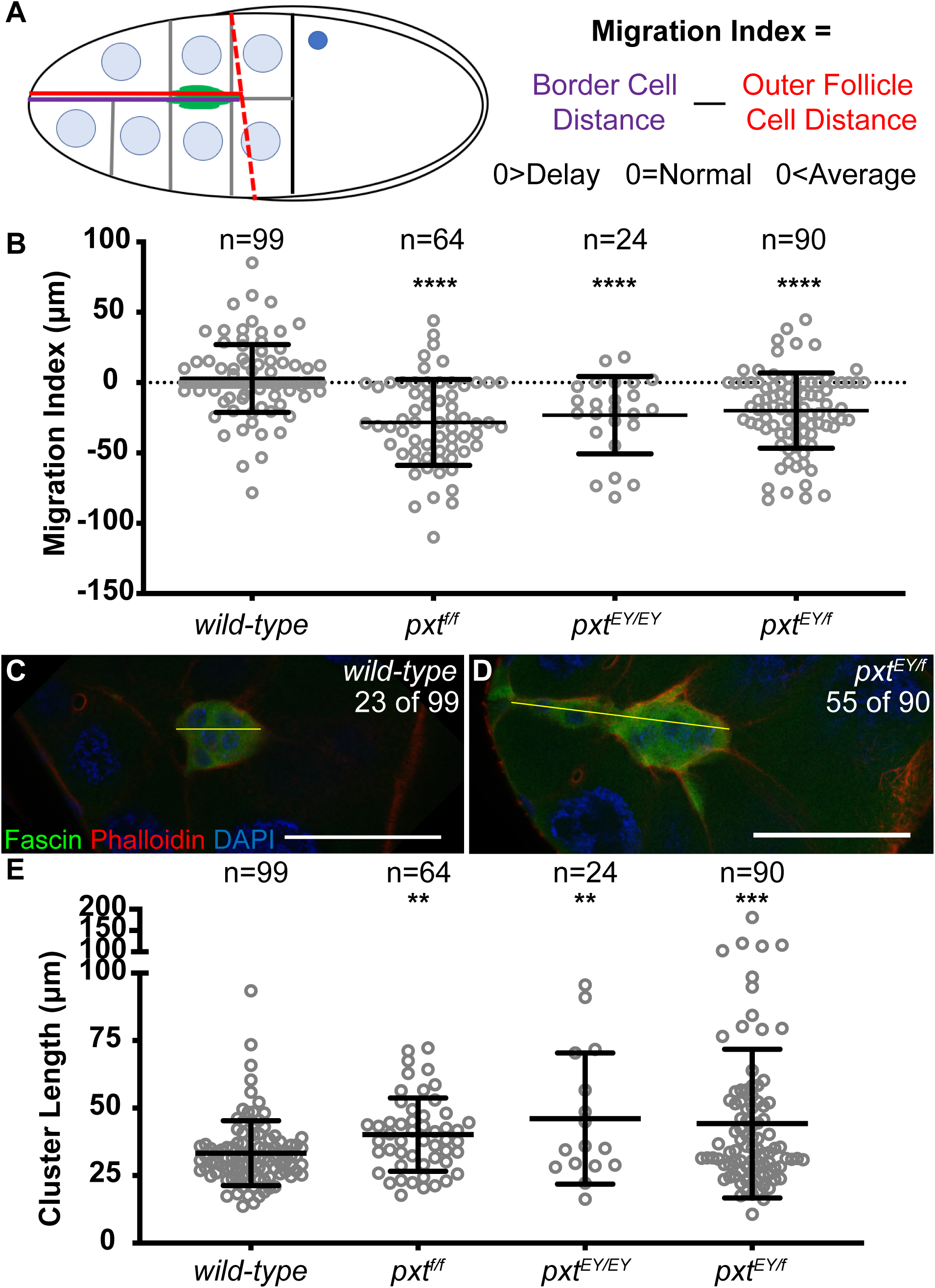
Prostaglandins regulate S9 border cell migration and cluster morphology. A. Diagram depicting the measurements used to quantify border cell migration during S9. The position of the border cells is quantified by measuring the distance from the anterior tip of the follicle to the leading edge of the border cell cluster (A, purple line), and the position of the outer follicle cells is quantified by measuring from the anterior tip of the follicle to the anterior edge of the outer follicle cells (A, red line; red dashed line indicates the position of the outer follicle cells). The difference between the border cell distance (A, purple) and the outer follicle cell distance is termed the migration index (A, red); units = μm. Normal or on-time migration should result in a migration index of 0, while delayed migration will result in negative values and accelerated migration will result in a positive value. B. Graph of the migration index quantification during S9 for the indicated genotypes. Each circle represents a single border cell cluster; the line indicates the average and the whiskers indicate the standard deviation (SD). Dotted line at 0 indicates an on-time migration. C-D. Maximum projection of 3 confocal slices of S9 follicles of the indicated genotypes; anterior is to the left. C. *wild-type* (*yw*). D. *pxt^EY/f^*. Merged images: Fascin, green; Phalloidin (F-actin), red; and DAPI (DNA), blue. The frequency of S9 follicles exhibiting rearward elongated border cell clusters is indicated at the top right of the panels. The yellow lines in C-D represent the measurements assessed in E. E. Graph of the quantification of primary cluster length for the indicated genotypes; note that cells left behind and fully detached from the cluster were not included in the measurements. Each circle represents a single border cell cluster; the line indicates the average and the whiskers indicate the SD. While wild-type follicles exhibit on-time border cell migration (B) and a round cluster morphology (C, E), loss of Pxt results in significantly delayed border cell migration and elongated clusters with trailing cells during S9 (D, E). ****p <0.0001, ***p<0.001, and **p<0.01. Scale bars = 50μm.

In addition to delayed migration, loss of Pxt also alters border cell cluster morphology during S9. Wild-type clusters are round and held tightly together with one main protrusion coming from the front of the cluster (Figure 2C, (Bianco *et al*., 2007; Prasad and Montell, 2007)). We find that ∼23% of wild-type clusters have a posterior protrusion or tail. However, in *pxt* mutants the majority of border cell clusters are elongated and have tails (61%, Figure 2D). To further quantify this defect, we measured the length of the clusters (Figure 3C and D yellow line) and found that compared to wild-type the loss of Pxt results in significantly longer clusters (Figure 3E). Wild type clusters averaged 33.32µm in length while clusters in *pxt^f/f^* follicles averaged 40.20µm (p=0.0018), clusters in *pxt^EY/EY^* follicles averaged 46.13µm (p=0.0012), and clusters in *pxt^f/EY^* follicles averaged 44.25µm (p=0.0004). These data reveal that Pxt regulates the morphology of the border cell cluster and suggest that the shape defects may impair migration. Furthermore, this change in cluster morphology likely accounts for the increased number of unattached cells observed at S10 in *pxt* mutants (Supplementary Figure 1A).

### Prostaglandin signaling is necessary in the somatic cells for on-time border cell migration

Having found that Pxt is required for border cell migration during S9, we next sought to determine where Pxt activity is necessary. Pxt is expressed in all cells of the developing follicle (Tootle and Spradling, 2008). Thus, Pxt may function in the germline cells, the somatic cells, or both to promote proper border cell migration.

To assess where Pxt is required we used the UAS/GAL4 system (Fischer *et al*., 1988) to knockdown Pxt by RNAi in the somatic cells (*c355 GAL4*). As expected the GAL4 only control exhibited normal border cell migration (Figure 4A). RNAi knockdown of Pxt in the somatic cells results in delayed border cell migration (Figure 4B). Quantification of the migration index reveals that somatic knockdown of Pxt resulted in a significant delay with a migration index of - 23.8µm compared to the control migration index of 4.34µm (Figure 4C, p=0.0005). This finding was verified using a second RNAi line (Supplementary Figure 4). Unfortunately, these RNAi constructs are under the control of the UASt promoter, which cannot be expressed with germline GAL4 drivers. Together these data indicate that Pxt is required in the somatic cells to regulated on-time border cell migration.

**Figure 4:**
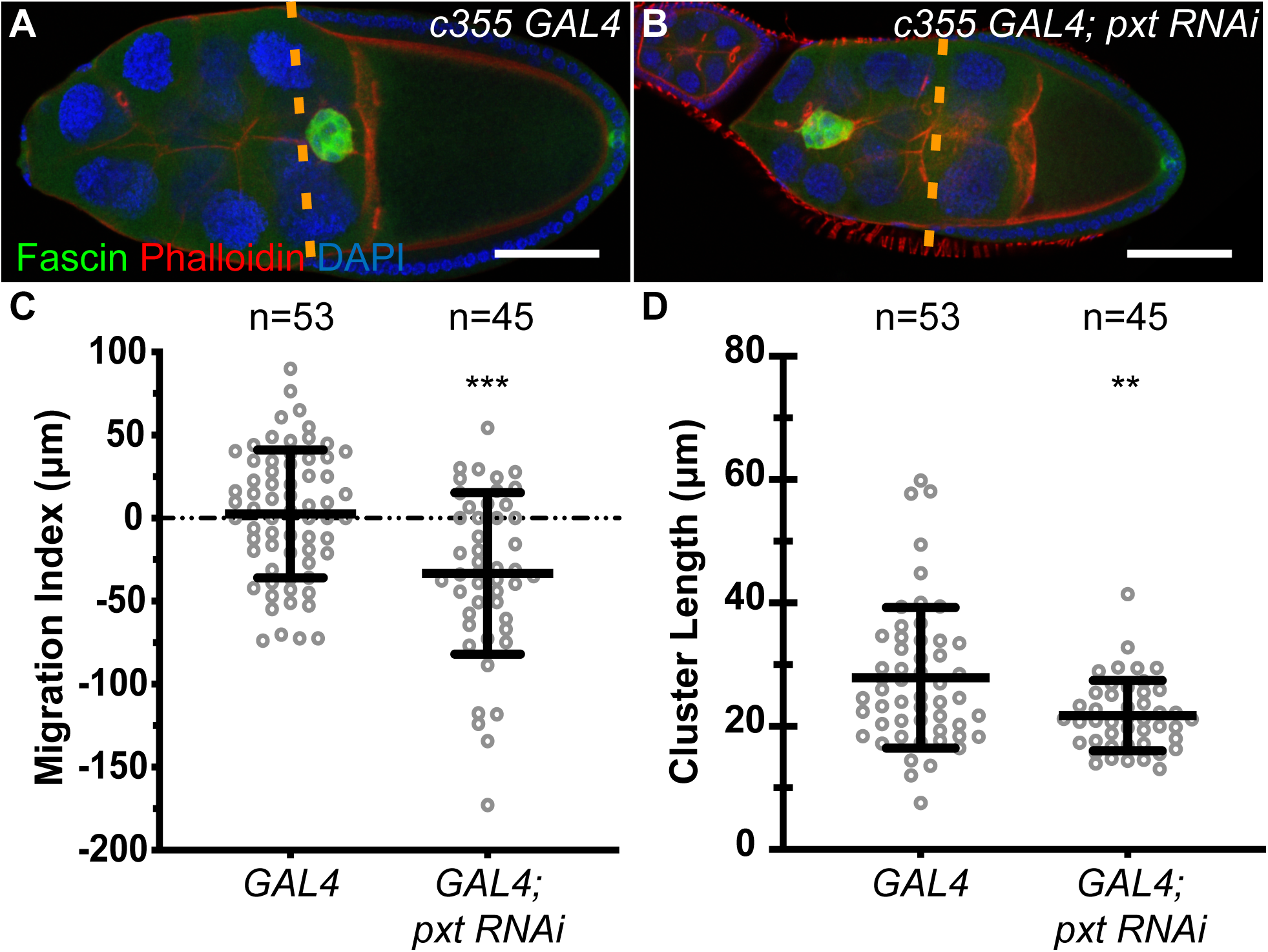
Pxt is required in the somatic cells for on-time border cell migration. A-B. Maximum projection of 3 confocal slices of S9 follicles of the indicated genotypes; anterior is to the left. A. Somatic GAL4 control (*c355 GAL4/+*). B. Somatic knockdown of Pxt (*c355 GAL4/+; pxt RNAi/+*). Merged images: Fascin, green; Phalloidin (F-actin), red; and DAPI (DNA), blue. Dashed orange lines indicate the position of the outer follicle cells. C. Graph of the migration index quantification during S9 for the above indicated genotypes. D. Graph of the quantification of primary cluster length for the above indicated genotypes; measured as described in Fig. 3. In C-D, each circle represents a single follicle; the line indicates the average and the whiskers indicate the SD. Somatic knockdown of Pxt results in delayed border cell migration (B, C) and shorter cluster length (D), compared to somatic GAL4 controls (A, C-D). ***p<0.001, and **p<0.01. Scale bars = 50μm.

We next assessed how somatic knockdown of Pxt affects cluster morphology. While qualitative analysis of the fixed images did not reveal striking cluster morphology defects when Pxt was knockdown (Fig. 4B compared to A), quantitative analysis uncovered a surprising result. Somatic knockdown of Pxt results in a more condensed cluster, with an average length of 21.73µm compared to 27.85µm for the control clusters (Figure 4D, p=0.0015). Interestingly this does not seem to be due to a change to either an increase in detached cells or due to a change in the number of cells in the cluster (Supplementary Figure 1C-D). These findings reveal that somatic knockdown of Pxt is not sufficient to cause the elongated cluster morphology observed in the *pxt* mutant follicles. This difference may be due to insufficient loss of Pxt by RNAi knockdown and/or that Pxt in the germline modulates cluster morphology. We favor the latter possibility.

### Loss of prostaglandin signaling alters integrin localization

Having found that loss of Pxt results in both delayed migration and aberrant, elongated clusters, we hypothesized that these defects could be the result of abnormal cellular adhesions. Two adhesion factors that regulate border cell migration are E-cadherin (Niewiadomska *et al*., 1999; De Graeve *et al*., 2012; Cai *et al*., 2014) and integrins (Dinkins *et al*., 2008; Llense and Martin-Blanco, 2008). Both increased and decreased E-cadherin levels in either the border cells or the nurse cells inhibit border cell migration (Niewiadomska *et al*., 1999; Cai *et al*., 2014). We find that E-cadherin localization and levels appears grossly normal in *pxt* mutants (Supplementary Figure 5). Integrin receptors are composed of one alpha and one beta subtype. In the border cells, βPS-integrin (*Drosophila* Myospheroid, Mys) and *α*PS3-integrin (*Drosophila* Scab, Scb) are enriched on the border cell membranes; RNAi knockdown either results in delayed border cell migration during S9 (Dinkins *et al*., 2008).

We find that while integrin appears normal in the outer follicle cells (data not shown), integrin localization is strikingly altered in the border cells of *pxt* mutants. To account for potential staining variability, we stained wild-type and *pxt* mutant follicles for βPS-integrin in the same tube. In wild-type follicles, βPS-integrin exhibits strong and continuous stretches of membrane localization on the border cell cluster (Figure 5A and C). Conversely, loss of Pxt results in variable integrin levels and localization. In general, when stained in the same tube as wild-type follicles, *pxt* mutant follicles exhibit reduced enrichment of integrin at the membrane and increased cytoplasmic integrin (Figure 5B and D). Indeed, line scan analysis of relative fluorescence intensity along the yellow dashed lines reveals that in wild-type follicles the border cells have high peaks of integrin intensity at the membrane (Figure 5A’ and C’, red asterisks), while in *pxt* mutant follicles the clusters lack membrane enrichment and have integrin staining throughout (Figure 5B’) or exhibit less membrane enrichment (Figure 5D’, red asterisk). To assess the frequency of these differences, follicles were scored as having high or low membrane staining and high or low cytoplasmic staining in a genotypically blinded manner (see Materials and Methods for details). We find that compared to their wild-type counterparts, follicles that have lost Pxt are more likely to have low membrane staining (73% compared to 38%) and high cytoplasmic staining (73% compared to 38%) (Figure 5E, p<0.0001). In Figure 5, both wild-type examples were scored as having high membrane and low cytoplasmic staining (Figure 5A, C), while the *pxt^EY/EY^* example has low membrane and high cytoplasmic staining (Figure 5B) and the *pxt^f/f^* example has low membrane and low cytoplasmic staining (Figure 5D). These data suggest that PGs are needed for the correct membrane enrichment of integrins on the border cells.

**Figure 5:**
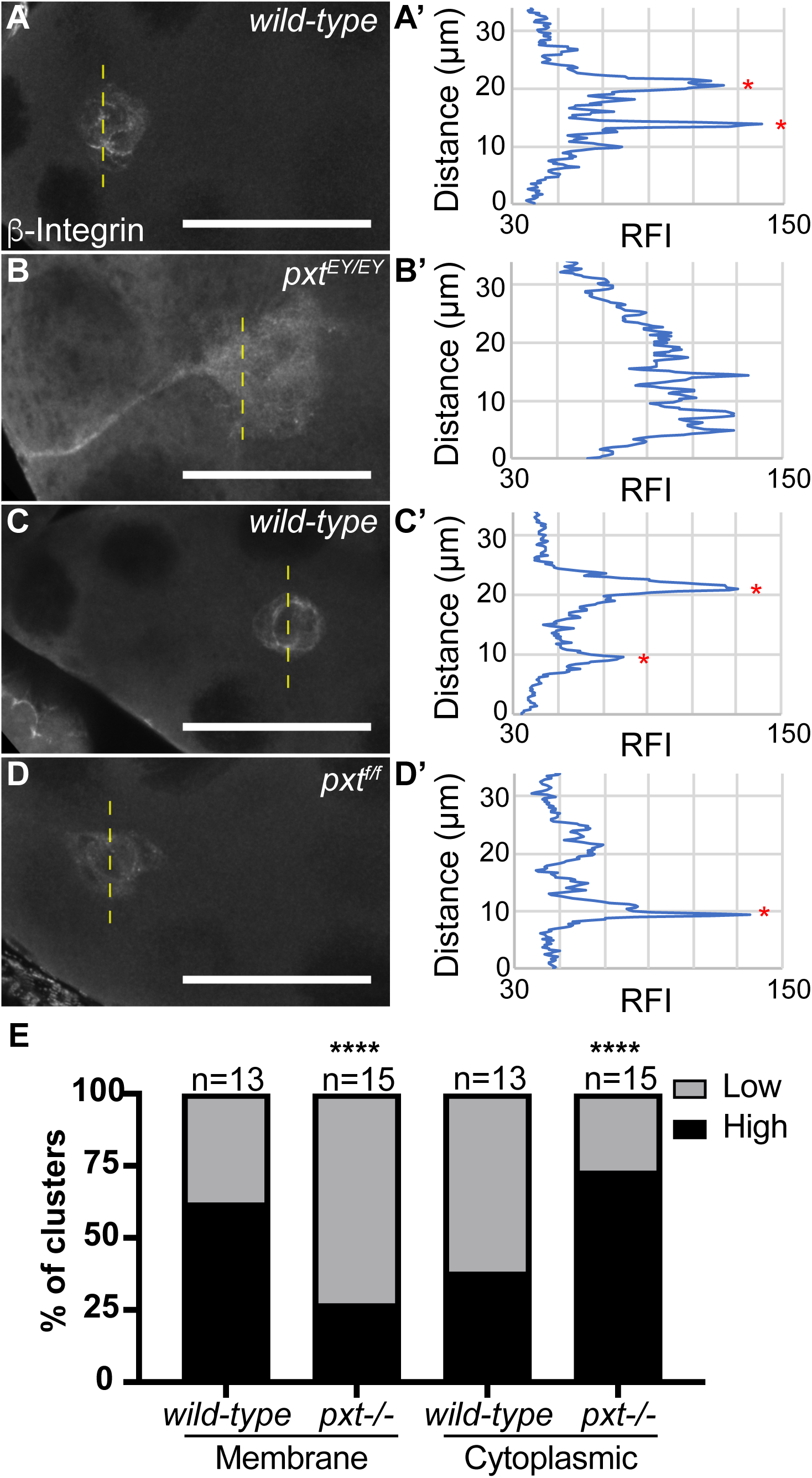
Pxt is required for integrin localization in the border cell cluster. A-D. Maximum projection of 3 confocal slices of S9 border cell clusters of the indicated genotypes stained with β-integrin (white); anterior is to the left. A, C. wild-type (*yw*). B. *pxt^EY/EY^*. D. *pxt^f/f^*. A’-D’. Graphs of the line scans of relative fluorescent intensity (RFI) across the yellow dashed lines in A-D, respectively. A-B’ and C-D’ represent paired images of wild-type and a *pxt* mutant at a similar point during the border cell migration. E. Graph showing the quantification of membrane and cytoplasmic β-integrin intensity in the border cell clusters of wild-type and *pxt* mutant follicles. In a genotypically blinded manner, confocal images of border cell clusters were scored as having either high (black) or low (grey) membrane staining and high (black) or low (grey) cytoplasmic staining. In wild-type border cell clusters there is bright localization of integrin to the membrane (A, C); this is evident in line scan graphs where peaks of membrane staining are marked with red asterisks (A’, C’) and from binning clusters into high and low membrane and cytoplasmic staining (E). In *pxt* mutants the membrane localization is reduced and there is higher cytoplasmic integrin levels (B-B’, D-D’, E). ****p<0.0001. Scale bars = 50μm.

To further assess the relationship between PGs and integrins, we utilized dominant genetic interactions. Heterozygosity for a mutation in either βPS-integrin (*mys^10^/+*), *α*PS3-integrin (*scb^01288^/+*), or *pxt* alone should have no effect on border cell migration. If the border cell migration defects in *pxt* mutants are due to reduced integrin, then double heterozygotes (*mys^10^/+; pxt-/+* or *scb^01288^/+; pxt-/+*) should exhibit migration defects. However, we find that co-reduction of Pxt and one integrin subunit exhibits normal border cell migration (Supplemental Figure 6). These findings may indicate that heterozygosity for an integrin subunit is not a sufficient enough reduction to observe an interaction, or that the integrin phenotype in *pxt* mutants is not reflective of a simple reduction in integrins. Instead, Pxt may regulate integrins by controlling their activation, and subsequent clustering and stabilization of the receptors on the membranes. We favor this latter possibility.

### Prostaglandins regulate on-time border cell migration and integrin localization through the actin bundling protein Fascin

We next wanted to determine how PGs regulate integrin localization in the border cell cluster. Integrins can be activated by extracellular matrix (ECM) ligands or intracellularly by the actin cytoskeleton (Harburger and Calderwood, 2009; Vicente-Manzanares *et al*., 2009). The latter is thought to be the primary means activating integrins in the border cells, as there is only one reported case of a puncta of ECM contributing to border cell migration (Medioni and Noselli, 2005). Previously, we found that one way PGs regulate the actin cytoskeleton during oogenesis is through promoting the activity of the actin bundling protein, Fascin (Groen *et al*., 2012). Fascin is highly expressed in the border cell cluster (Cant *et al*., 1994) and we find that loss of Fascin causes delayed border cell migration during S9 (Lamb *et al*., 2019). Therefore, we hypothesized that PGs regulate Fascin to control integrins in the border cell cluster.

If our hypothesis is correct, then loss of Fascin is predicted to alter integrin levels and localization. We stained wild-type and *fascin*-null follicles for βPS-integrin in the same tube to account for potential staining variability. The *fascin*-null clusters display altered integrin localization similar to the *pxt* mutants, with low membrane localization and high cytoplasmic staining (Fig. 6A, B). Using the same quantification method described above of scoring high or low localization of integrin in a genotypically double blinded manner, we find the *fascin*-null clusters have a significant increase in frequency of clusters with high cytoplasmic staining and a decreased frequency of high membrane staining compared to the wild-type controls (Fig. 6C, p<0.0001). In Figure 6, both wild-type examples were scored as having high membrane and low cytoplasmic integrin staining (Figure 6A-B), while one *fascin* mutant example has high membrane and high cytoplasmic staining (Figure 6C) and the other has low membrane and high cytoplasmic staining (Figure 6D). The similar phenotypes in the *fascin* and *pxt* mutants support the model that they act together to regulate integrins.

**Figure 6:**
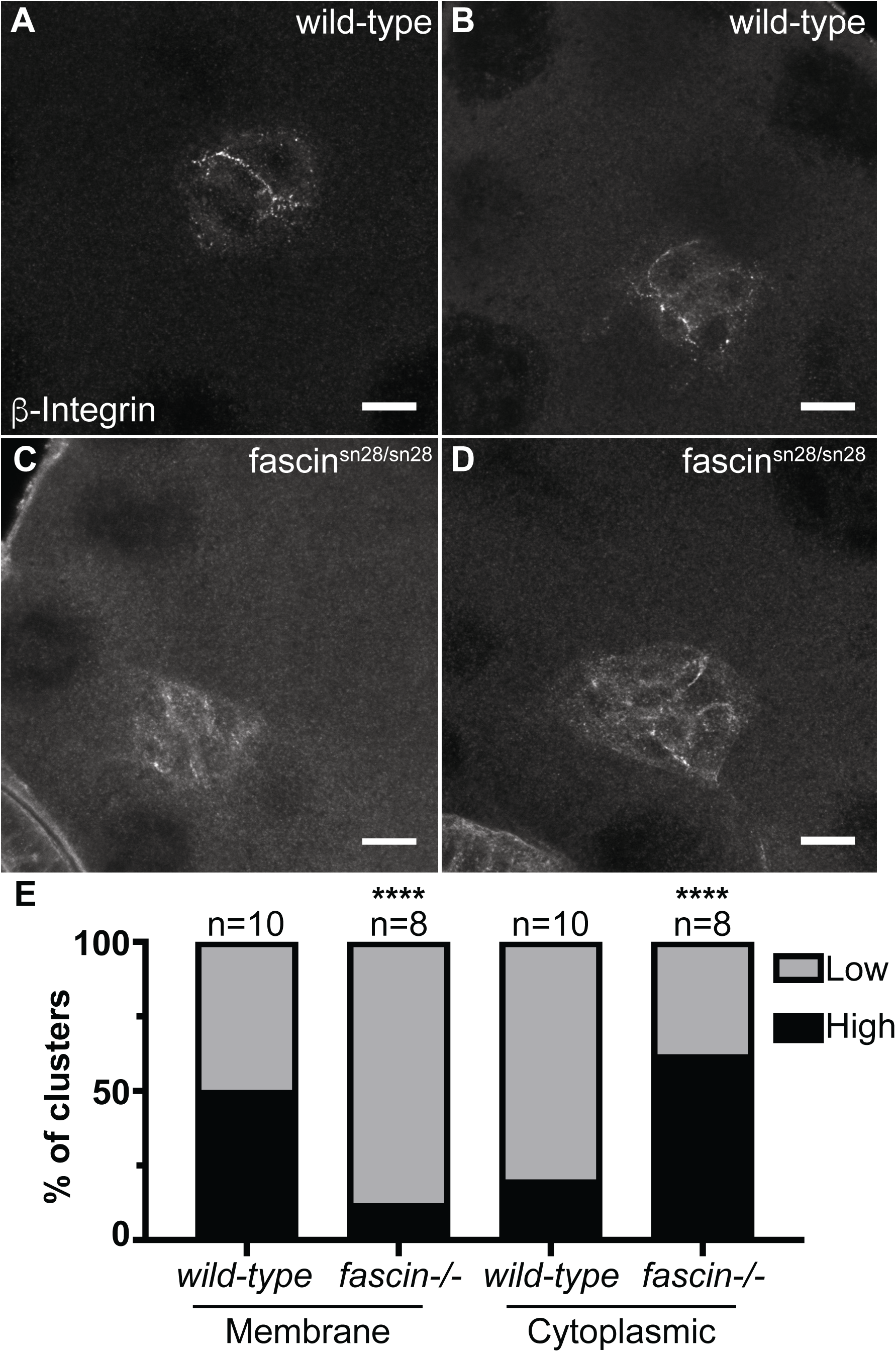
Fascin is essential for proper integrin localization in the border cell cluster. A-D. Maximum projections of 3 confocal slices of S9 border cell clusters of the indicated genotypes stained with β-integrin (white); anterior is to the left. For ease of visualization, all images brightened by 50% in Photoshop. A-B. wild-type C-D. *fascin^sn28/sn28^*. E. Graph showing quantification of membrane and cytoplasmic β-integrin intensity in the border cells of wild-type and *fascin*-null follicles for two independent experiments. Clusters were scored in a double blinded manner for whether they had high (black) or low (grey) membrane and cytoplasmic intensity. Loss of Fascin results in altered integrin localization in the border cell cluster. Similar to *pxt* mutants, the clusters in *fascin*-null follicles display a higher frequency of low membrane intensity and high cytoplasmic intensity compare to *wild-type* controls (C-D, compared to A-B, and E). ****p<0.001. Scale bars= 10μm.

To further test our hypothesis, we used dominant genetic interaction studies to determine if PGs regulate Fascin to promote border cell migration. Partial reduction of either Pxt (*pxt+/-*) or Fascin (*fascin+/-*) should not alter border cell migration. However, if PGs and Fascin function together to promote border cell migration, then reduced levels of both (*fascin+/-; pxt+/-*) will display defects in border cell migration. We performed immunofluorescent staining for Hts and FasIII; this stain labels both the border cells and outer follicle cells and enables us to assess border cell migration in a similar manner to the Fascin stain used previously. As expected, heterozygous loss of Pxt or Fascin alone does not alter border cell migration using our migration index quantification (Figure 7C). However, partial loss of both Pxt and Fascin (*fascin+/-; pxt+/-*) results in significant border cell migration delays (Figure 7A-C, average migration indices: 25.47 for *fascin^sn28^/+; pxt^EY^/+* compared to −4.58 for *pxt^EY^/+*, p=0.002; and −13.59 for *fascin^sn28^/+; pxt^f^/+* compared to 3.49 for *pxt^f^/+*, p=0.039). Additionally, we observed altered border cell cluster morphology in follicles heterozygous for mutations in both *pxt* and *fascin*. Similar to the clusters in *pxt* mutant follicles, the clusters from double heterozygous follicles (*fascin+/-; pxt+/-*) display an elongated phenotype with posterior tails (Figure 7D-E). Quantification reveals that the clusters from *fascin+/; pxt+/-* follicles have a significant increase in cluster length (Figure F; average cluster length: 39.69 for *fascin^sn28^/+; pxt^EY^/+* compared to 28.23 for *pxt^EY^/+*, p=0.0135; and 47.62 for *fascin^sn28^/+; pxt^f^/+* compared to 27.71 for *pxt^f^/+*, p<0.0001). These results indicate that PGs and Fascin genetically interact to regulate border cell migration and morphology.

**Figure 7:**
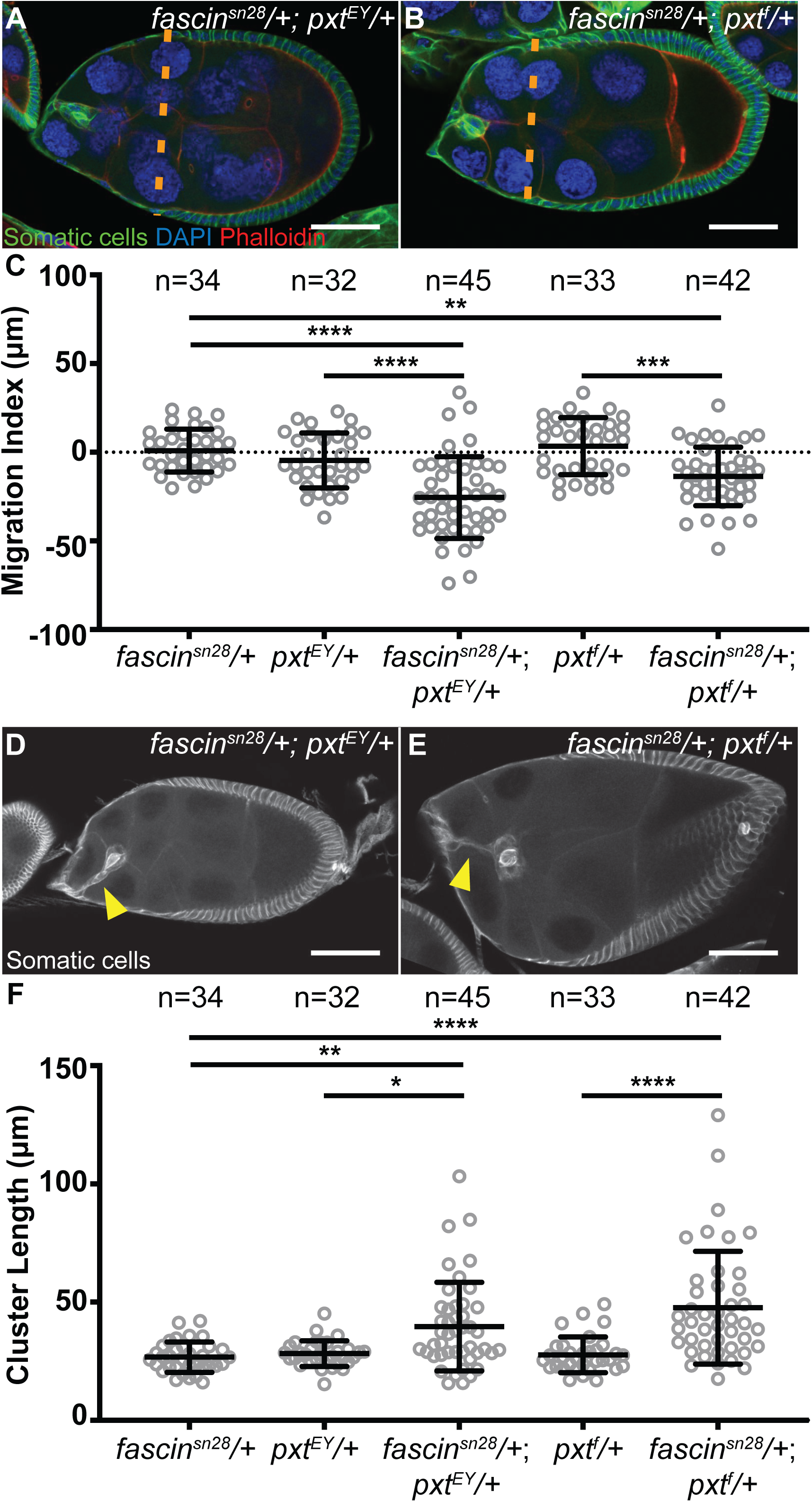
Prostaglandins regulate Fascin to promote border cell migration. A-B. Maximum projections of 2-4 confocal slices of S9 follicles of the indicated genotypes; anterior is to the left. A. *fascin^sn28^/+; pxt^EY^/+*. B. *fascin^sn28^/+; pxt^f^/+*. Merged images: DAPI (blue), Phalloidin (red), and Somatic Cell stain (Hts and FasIII, green). Orange dashed lines indicate the position of the outer follicle cells. C. Graph of the migration index quantification during S9 border cell migration in the indicated genotypes. Each circle represents a single border cell cluster; the line indicates the average and the whiskers indicate the SD. D-E. Maximum projections of 2-4 confocal slices of S9 follicles of the indicated genotypes stained with the somatic cell stain (Hts and FasIII, white). D. *fascin^sn28^/+; pxt^EY^/+*. E. *fascin^sn28^/+; pxt^f^/+*. Yellow arrowheads denote tails attached to the cluster. F. Graph of the quantification of border cluster length from follicles of the indicated genotypes. Each circle represents a single border cell cluster; the line indicates the average and the whiskers indicate the SD. While heterozygosity for mutations in *pxt* or *fascin* do not affect border cell migration or cluster morphology, double heterozygotes (*fascin-/+;pxt-/+*) exhibit delayed border cell migration (A-C) and elongated clusters (D-F). ****p<0.0001, ***p<0.001, **p<0.01 and *p<0.05. Scale bars= 50μm.

## Discussion

*Drosophila* border cell migration has been widely used to identify factors regulating invasive, collective cell migration (Montell, 2003; Montell *et al*., 2012). Here we find that PG signaling is required for on-time border cell migration and normal cluster morphology. Specifically, loss of the COX-like enzyme Pxt results in both delayed border cell migration during S9 and aberrant, elongated clusters, with cells occasionally being left behind along the migration path.

The border cell defects vary in severity across the two *pxt* alleles – *pxt^f^* and *pxt^EY^*. Both alleles result in delayed migration and increased cluster length during S9 (Figures 2-3). During S10, the stronger allele, *pxt^f^*, results in elongated clusters with too many cells, while the weaker allele, *pxt^EY^*, has a lower frequency of elongated clusters (Figure 1 and Supplemental Figure 1). Transalleles of *pxt^EY/f^* exhibit an intermediate phenotype. These data suggest that the phenotypic variation is primarily due to the level of Pxt loss.

RNAi knockdown of Pxt in the somatic cells only partially recapitulates the defects observed in *pxt* mutants. Somatic knockdown of Pxt causes delayed border cell migration during S9. However, the border cell clusters are not elongated, but instead exhibit a shorter length (Figure 4). The phenotypic differences between the *pxt* mutants and RNAi knockdown may be due to the RNAi failing to reduce Pxt sufficiently. However, if this were the case, the cluster length should be normal and not shorter. These data lead us to hypothesize that Pxt acts in both the somatic and the germ cells to regulate border cell migration. Specifically, Pxt may function within the somatic cells, likely within the border cells themselves, to mediate on-time border cell migration. While Pxt may function within the germ cells, likely within the nurse cells on which the border cells migrate, to control cluster morphology. The germline role of Pxt could be to produce PGs that signal to the border cells to regulate cluster cohesion. Alternatively, PGs may act within the nurse cells to control their stiffness and/or adhesion to each other, and thereby, affecting border cell morphology. Indeed, the balance of force between the border cell cluster and the nurse cells must be maintained in order for normal cluster morphology, and misbalanced forces in either tissue lead to border cell migration delays and cluster elongation (Aranjuez *et al*., 2016). Supporting a role for PGs in regulating nurse cell stiffness, *pxt* mutants exhibit altered levels and localization of phospho-myosin regulatory light chain (*Drosophila* Spaghetti Squash) on the nurse cells during S10B (Spracklen *et al*., 2019); a known regulator of nurse cell stiffness (Aranjuez *et al*., 2016). Additionally, loss of PGs also causes cortical actin defects and breakdown (Tootle and Spradling, 2008; Groen *et al*., 2012). Together these data lead us to speculate that within the nurse cells PGs act to maintain proper cortical actin and stiffness, and in the absence of Pxt, the nurse cells are softer, resulting in border cell cluster morphology changes.

The idea that PGs regulate cell migration within both the migrating cells and their microenvironment is conserved across organisms. Cancer cells upregulate COX enzyme expression and exhibit increased PG production (Cha and DuBois, 2007; Wang and Dubois, 2010; Menter and Dubois, 2012). Specifically, increased PG signaling is associated with increased *in vitro* cell migration and invasion that can be blocked by COX inhibitor treatment (Tsujii *et al*., 1997; Chen *et al*., 2001; Lyons *et al*., 2011). Increased PG production within the tumor cells is also associated with high levels of *in vivo* metastasis and poor patient outcomes (Rolland *et al*., 1980; Khuri *et al*., 2001; Gallo *et al*., 2002; Denkert *et al*., 2003). These data support a role for PGs within the migrating cells themselves. PGs also play a role in the tumor microenvironment, contributing to chronic inflammation and immune modulation (Wang and DuBois, 2018). Additionally, in a key study, Li et al. 2012 uncovered that PG signaling within the mesenchymal stroma cells play critical roles in regulating the fate of the carcinoma cells and promoting the cells to undergo an epithelial to mesenchymal transition and invade the surrounding tissue (Li *et al*., 2012). Thus, it is critical to define how PGs act both within the migrating cells and their environment to control invasion.

One means by which PGs regulate migration is through modulating integrins. Integrins are cell adhesion receptors that can be activated by binding to ECM components (termed outside-in signaling) or can be activated by intracellular changes in the actin cytoskeleton connecting to the intracellular domains of the integrins (termed inside-out signaling; (Harburger and Calderwood, 2009; Vicente-Manzanares *et al*., 2009)). In the context of mammalian cultured cells, high COX levels and PGE_2_ production are associated with increased integrin expression and accumulation on the cell surface (Mayoral *et al*., 2005; Liu *et al*., 2010). PGF_2_*_α_*-mediated cell adhesion is inhibited by blocking integrin activity (Sales *et al*., 2008). Additionally, COX inhibition blocks signaling downstream of integrin receptors (Dormond *et al*., 2001; Dormond *et al*., 2002). Supporting that PG regulation of integrin signaling is conserved in *Drosophila*, we find that loss of Pxt results in decreased enrichment of high levels of βPS-integrin on the border cell membranes (Figure 5).

During border cell migration, integrin signaling is poorly understood. There is little evidence of ECM surrounding the border cells or contributing to their migration (Medioni and Noselli, 2005). However, integrins are enriched on the border cell membranes (Figure 5; (Dinkins *et al*., 2008)). Additionally, RNAi knockdown of βPS-integrin or *α*PS3-integrin results in delayed border cell migration during S9 (Dinkins *et al*., 2008). These data suggest that integrins have a role in mediating on-time border cell migration and that they are likely activated by actin cytoskeletal changes.

Previously, we established the PGs regulate actin dynamics necessary for late-stage *Drosophila* follicle morphogenesis by controlling a number of actin binding proteins, including the actin bundler Fascin (Groen *et al*., 2012; Spracklen *et al*., 2014; Spracklen *et al*., 2019). Fascin is highly upregulated in the border cells (Cant *et al*., 1994) and we recently found that Fascin is required for on-time border cell migration during S9 (Lamb *et al*., 2019). These data led us to hypothesize that within the border cells PGs may regulate Fascin and thereby, the actin cytoskeleton. The cytoskeletal changes in turn modulate integrin localization, and together these factors mediate on-time border cell migration and cluster morphology. Supporting this hypothesis loss of Fascin results in *pxt*-like defects in βPS-integrin levels and localization on the border cell cluster (Figure 6). Further, dominant genetic interaction studies reveal that S9 follicles from double heterozygotes, *fascin-/+; pxt-/+*, phenocopy *pxt* mutants. Specifically, border cell migration is delayed during S9 and the clusters are elongated (Figure 7). These data indicate that PGs regulate Fascin to control both border cell migration and cluster morphology.

As both Pxt and Fascin regulate integrin enrichment on the border cell cluster, we hypothesize this contributes to border cell migration and morphology. The decreased membrane enrichment of integrins in either *pxt* or *fascin* mutants could be caused by a number of mechanisms. Two potential causes are decreased expression or decreased trafficking of integrins to the cell surface. If either of these were the case, then the dominant genetic interactions between mutations in the integrin subunits and *pxt* would be expected to cause border cell migration defects. Instead, we find that migration is normal (Supplemental Figure 6). Additionally, our prior microarray analysis indicates that PGs do not alter integrin expression during *Drosophila* oogenesis (Tootle *et al*., 2011). Another means of affecting integrin enrichment is decreased activation and thus, decreased clustering. Both activation and clustering of integrins require interaction with the actin cytoskeleton and adaptor proteins, including Paxillin (Harburger and Calderwood, 2009; Vicente-Manzanares *et al*., 2009). Notably, microarray analysis of *pxt* mutant follicles revealed that *paxillin* is downregulated (Tootle *et al*., 2011). Further, in mammalian systems, both PGs (Mayoral *et al*., 2005; Bai *et al*., 2009; Liu *et al*., 2010; Bai *et al*., 2013) and Fascin (Anilkumar *et al*., 2003; Villari *et al*., 2015) mediate increased integrin adhesion stability. These data lead us to speculate that the decreased membrane enrichment of integrins in both *fascin* and *pxt* mutants is due to a loss of inside-out activation of the integrins.

In conclusion, this study has uncovered a new pathway by which PGs regulate Fascin to regulate collective, invasive cell migration. One downstream effect of this pathway is altered integrin localization, and future studies are needed to elucidate the precise mechanisms involved. Additionally, we postulate that other Fascin-dependent activities are also controlled by PG signaling to mediate cluster invasion (Anilkumar *et al*., 2003; Villari *et al*., 2015; Jayo *et al*., 2016). Thus, border cell migration provides a robust, *in vivo* system to delineate the means by which PGs regulate Fascin-dependent collective, invasive cell migration. These same mechanisms likely contribute to cancer metastasis as both PGs (Rolland *et al*., 1980; Khuri *et al*., 2001; Gallo *et al*., 2002; Denkert *et al*., 2003) and Fascin (Hashimoto *et al*., 2004; Yoder *et al*., 2005; Okada *et al*., 2007; Li *et al*., 2008; Chan *et al*., 2010) are associated with highly aggressive cancers and poor patient outcomes.

## Materials and Methods

### Fly stocks

Fly stocks were maintained on cornmeal/agar/yeast food at 21°C, except where noted. Before immunofluorescence, flies were fed wet yeast paste daily for 2–4 d. *yw* was used as the *wild-type* control. The following stocks were obtained from the Bloomington Drosophila Stock Center (Bloomington, IN): *pxt^EY03052^* (BL15620), *c355 GAL4* (BL3750), *mys^10^* (BL58806), and *scb^01288^* (BL11035). The *pxt^f01000^* stock was obtained from the Harvard Exelixis Collection. The *UASt pxt RNAi* (V14379) and *UASt pxt RNAi* (V104446) stocks were obtained from the Vienna Drosophila Resource Center. The *sn28* line was a generous gift form Jennifer Zanet (Université de Toulouse, Toulouse, France; (Zanet *et al*., 2012)). Expression of the *UAS pxt RNAi* lines was achieved by crossing to the *c355* GAL4 line at room temperature and maintaining the adult progeny at 29°C for 3-5 days.

### Immunofluorescence

Whole-mount *Drosophila* ovary samples were dissected into Grace’s insect medium (Lonza, Walkersville, MD or Thermo Fischer Scientific, Waltham, MA) and fixed for 10 min at room temperature in 4% paraformaldehyde in Grace’s insect medium. Briefly, samples were blocked by washing in antibody wash (1X phosphate-buffered saline [PBS], 0.1% Triton X-100, and 0.1% bovine serum albumin) six times for 10 min each at room temperature. Primary antibodies were incubated overnight at 4°C, except for βPS-Integrin which was incubated for ∼20-48 h at 4°C. The following primary antibodies were obtained from the Developmental Studies Hybridoma Bank (DSHB) developed under the auspices of the National Institute of Child Health and Human Development and maintained by the Department of Biology, University of Iowa (Iowa City, IA): mouse anti-Fascin 1:25 (sn7c) (Cant *et al*., 1994); mouse anti-βPS-Integrin 1:10 (CF.6G11) (Brower *et al*., 1984); mouse anti-EYA 1:100 (eya10H6) (Boyle *et al*., 1997); mouse anti-β-catenin (N2 7A1, *Drosophila* Armadillo) 1:100 (Riggleman *et al*., 1990); rat anti-DCAD2 1:10 (Oda *et al*., 1994); mouse anti-Hts 1:50 (1B1) (Zaccai and Lipshitz, 1996); and mouse anti-FasIII 1:50 (7G10) (Patel *et al*., 1987)). Additionally, the following primary antibody was used, rabbit anti-Pxt 1:10000 (preabsorbed on *pxt^f/f^* ovaries at 1:20 and used at 1:500; (Spracklen *et al*., 2014)). After six washes in Triton antibody wash (10 min each), secondary antibodies were incubated overnight at 4°C or for ∼4 h at room temperature. The following secondary antibodies were used at 1:500–1:1000: AF488::goat anti-mouse, AF568::goat anti-mouse, AF647::goat anti-mouse, AF488::goat anti-rabbit, AF647::donkey anti-rabbit AF488::donkey anti-rat, and AF633::goat anti-rabbit (Thermo Fischer Scientific). Alexa Fluor 647–, rhodamine- or Alexa Fluor 488–conjugated phalloidin (Thermo Fischer Scientific) was included with secondary antibodies at a concentration of 1:100– 1:250. After six washes in antibody wash (10 min each), 4′,6-diamidino-2-phenylindole (DAPI, 5 mg/ml) staining was performed at a concentration of 1:5000 in 1X PBS for 10 min at room temperature. Ovaries were mounted in 1 mg/ml phenylenediamine in 50% glycerol, pH 9 (Platt and Michael, 1983). All experiments were performed a minimum of three independent times.

### Image acquisition and processing

Microscope images of fixed *Drosophila* follicles were obtained using LAS AF SPE Core software on a Leica TCS SPE mounted on a Leica DM2500 using an ACS APO 20×/0.60 IMM CORR -/D (Leica Microsystems, Buffalo Grove, IL), Zen software on a Zeiss 880 mounted on Zeiss Axio Observer.Z1 using Plan-Apochromat 20x/0.8 working distance (WD) = 0.55 M27, Plan-Apochromat 40x/1.3 Oil Differential Interference Contrast (DIC) WD = 2.0 or Plan-Apochromat 63x/1.4 Oil DIC f/ELYRA objectives, or Zen software on a Zeiss 700 LSM mounted on an Axio Observer.Z1 using a LD C-APO 40×/1.1 W/0 objective (Carl Zeiss Microscopy, Thornwood, NY). Maximum projections (two to five confocal slices), merged images, rotation, cropping, and distance measurements were performed using ImageJ software (Abramoff *et al*., 2004). *β*-integrin images on wild-type and *fascin* mutant clusters were all brightened by 50% in Photoshop to aid in visualization.

### Image Analyses

All image quantification was performed in a genotypically blinded manner, and where noted, in a double blinded manner, and was performed on a minimum of 3 independent experiments, except where noted in the Figure Legends.

Analysis of S10 clusters was performed on fixed confocal stacks of S10 follicles by counting the number of Eya stained nuclei visible within the border cell cluster and those left between the nurse cells using ImageJ software (Abramoff *et al*., 2004). The data was compiled, graphs generated and statistical analysis (Student’s t-test) were performed using GraphPad Prism version 7 or 8 (GraphPad Software, La Jolla California USA). In the graphs, the measurement for each follicle is represented as a circle, the averages and standard deviations are indicated by lines and whiskers, respectively.

To assess border cell migration during S9 a number of measurements were performed on fixed confocal stacks using Image J software (Abramoff *et al*., 2004). Specifically, we measured the follicle length, the distance between the anterior tip of the follicle and the leading edge (posterior) of the border cell cluster (distance of the border cells), the distance between the anterior tip of the follicle and the anterior edge of the outer follicle cells (distance of the follicle cells), and the distance from the rear to the front of the border cell cluster (cluster length; detached cells were not included in the length measurement). The data was compiled, and the migration index was calculated in Microsoft Excel (Microsoft, Redmond, WA). The migration index = distance of the outer follicle cells – distance of the border cells; units = μm. The migration index and cluster length data were compiled, graphs generated, and statistical analysis (Student’s t-test) were performed using GraphPad Prism version 7 or 8 (GraphPad Software). In the graphs, the measurement for each follicle is represented as a circle, the averages and standard deviations are indicated by lines and whiskers, respectively. To verify that any migration indices changes were due to altered border cell migration and not altered outer follicle cell position, the follicle length versus the distance of the outer follicle cells was plotted and analyzed in Microsoft Excel (Supplemental Figure 3).

Integrin localization and intensity were assessed by taking line scans of the relative fluorescent intensity across the border cell cluster of maximum projections of 3 slices of fixed, confocal stacks using ImageJ software (Abramoff *et al*., 2004). The line scan data was compiled and graphed in Microsoft Excel. Additionally, the integrin localization and intensity were analyzed from maximum projections of 3 slices of fixed, confocal stacks in a genotypically blinded manner or double blinded manner (see Figure Legends) by binning each cluster as having high or low membrane staining, and high or low cytoplasmic. Specifically, high membrane staining was defined as the cluster having more than one bright and continuous patches of integrin staining along the membrane; all non-high clusters were scored as having low membrane staining. High cytoplasmic staining was defined as having a high haze (at a similar intensity level of normal membrane staining in wild-type follicles) of integrin staining throughout the cytoplasm; all non-high clusters were scored as having low membrane staining. The data was compiled, graphs generated and statistical analysis (two-sided Fisher’s exact test) performed in GraphPad Prism version 7 or 8 (GraphPad Software).

## Acknowledgements

We thank the Westside Fly Group and Dunnwald lab for helpful discussions, and the Tootle lab for helpful discussions and careful review of the manuscript. Stocks obtained from the Bloomington Drosophila Stock Center (NIH P40OD018537) were used in this study. Information Technology Services – Research Services provides data storage support. This project is supported by National Institutes of Health R01GM116885. E.F.F has been supported by the NIH Predoctoral Training Grant in Genetics T32GM008629 (PI Daniel Eberl), the University of Iowa Graduate College Post-Comps Fellowship and Ballard and Seashore Fellowship. M.C.L. has been supported by the University of Iowa Summer Graduate Fellowship and the Anatomy and Cell Biology Department Graduate Fellowship. S.Q.M is supported by the NIH Predoctoral Training Grant in Genetics T32GM008629 (PI Daniel Eberl).

---

**Supplementary Figure 1:**
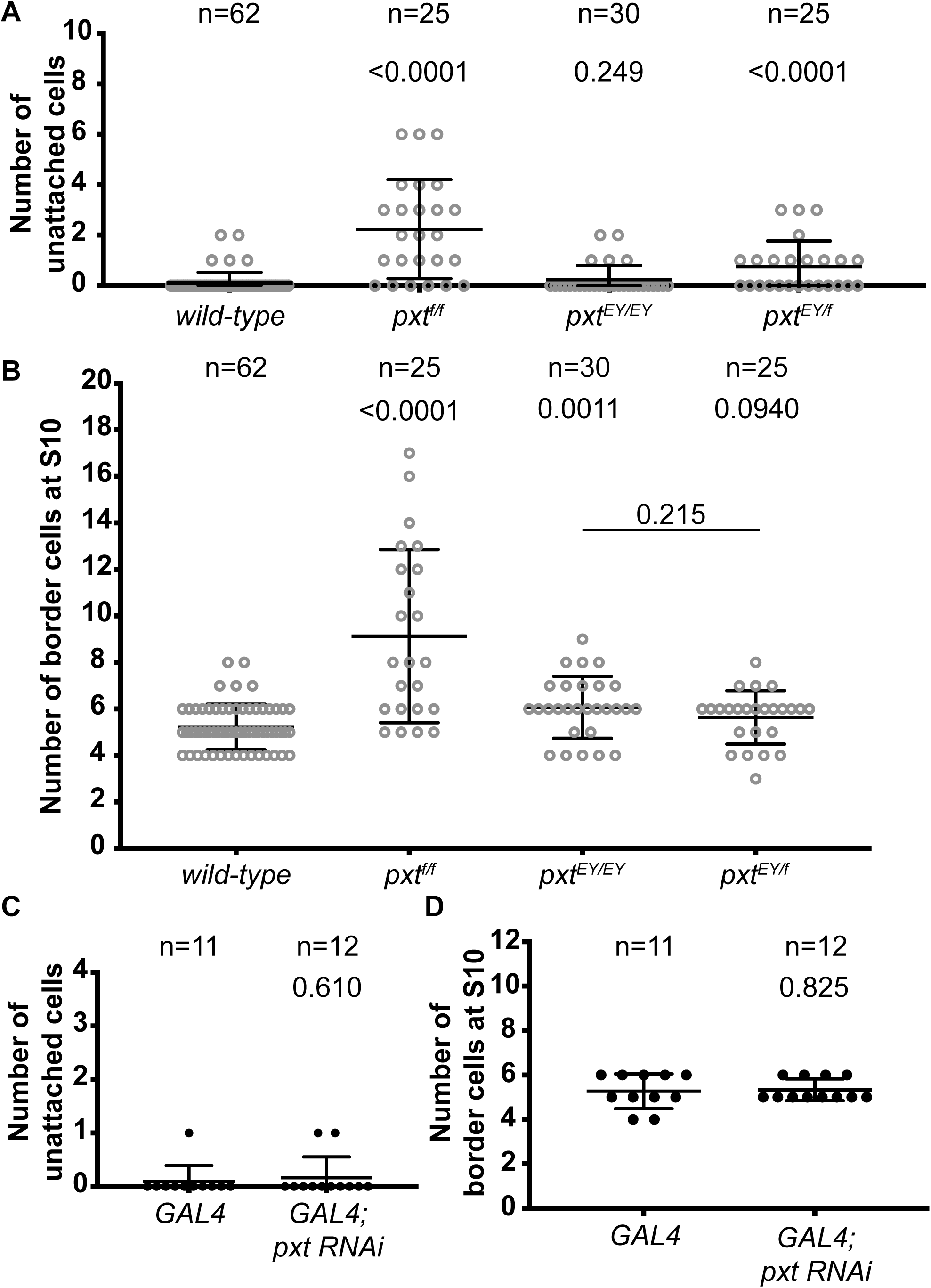
Pxt regulates border cell cluster morphology during S10. A, C. Graph of the quantification of the number of Eya stained somatic cells left between the nurse cells and visible during Stage 10 for the indicated genotypes. B, D. Graph of the quantification of the number of cells *pxt^EY/f^* within each border cell cluster, including trailing cells, at S10 for the indicated genotypes using Eya to mark the nuclei of the cells. In A-D, each circle represents a single border cell cluster; the line indicates the average and the whiskers indicate the SD. Loss of Pxt, via *pxt^f/f^* or *pxt^EY/f^*, results in border cells detaching and being left along the migration path (A). Additionally, *pxt^f/f^* results in an increase in the number of border cells (B). Conversely, somatic knockdown of Pxt does not results in border cells being left behind (C) or increased border cell number (D).

**Supplementary Figure 2:**
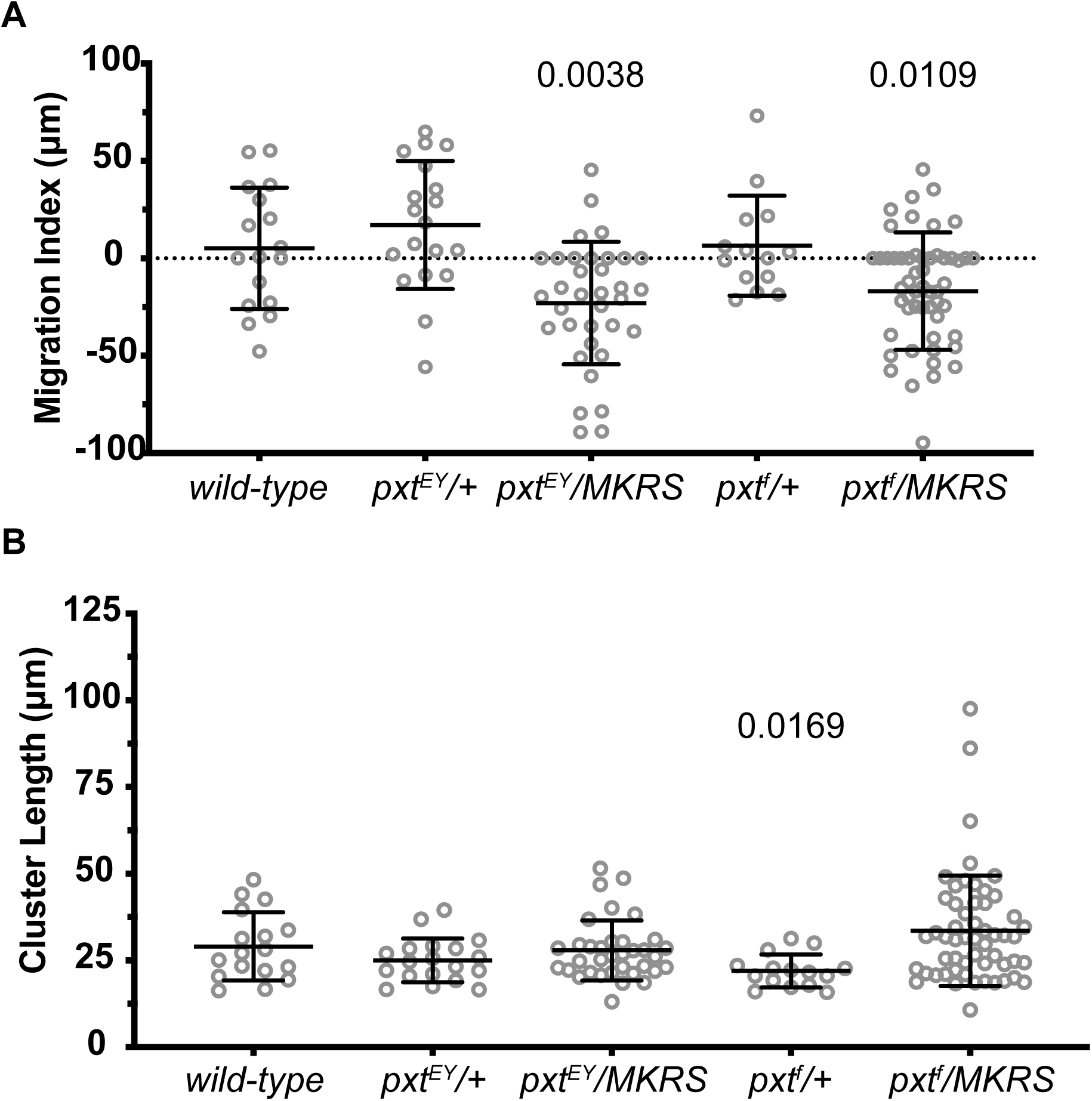
MKRS balancer genetically interacts with *pxt* mutants to cause border cell migration defects during S9. A. Graph of the migration index quantification during S9 for the indicated genotypes. B. Graph of the quantification of primary cluster length for the indicated genotypes; measured as described in Fig. 3. In A-B, each circle represents a single border cell cluster; the line indicates the average and the whiskers indicate the SD. Heterozygosity for *pxt* mutations over a wild-type chromosome has no effect on border cell migration or cluster length, while heterozygosity for a *pxt* mutation over the MKRS balancer results in delayed border cell migration.

**Supplementary Figure 3:**
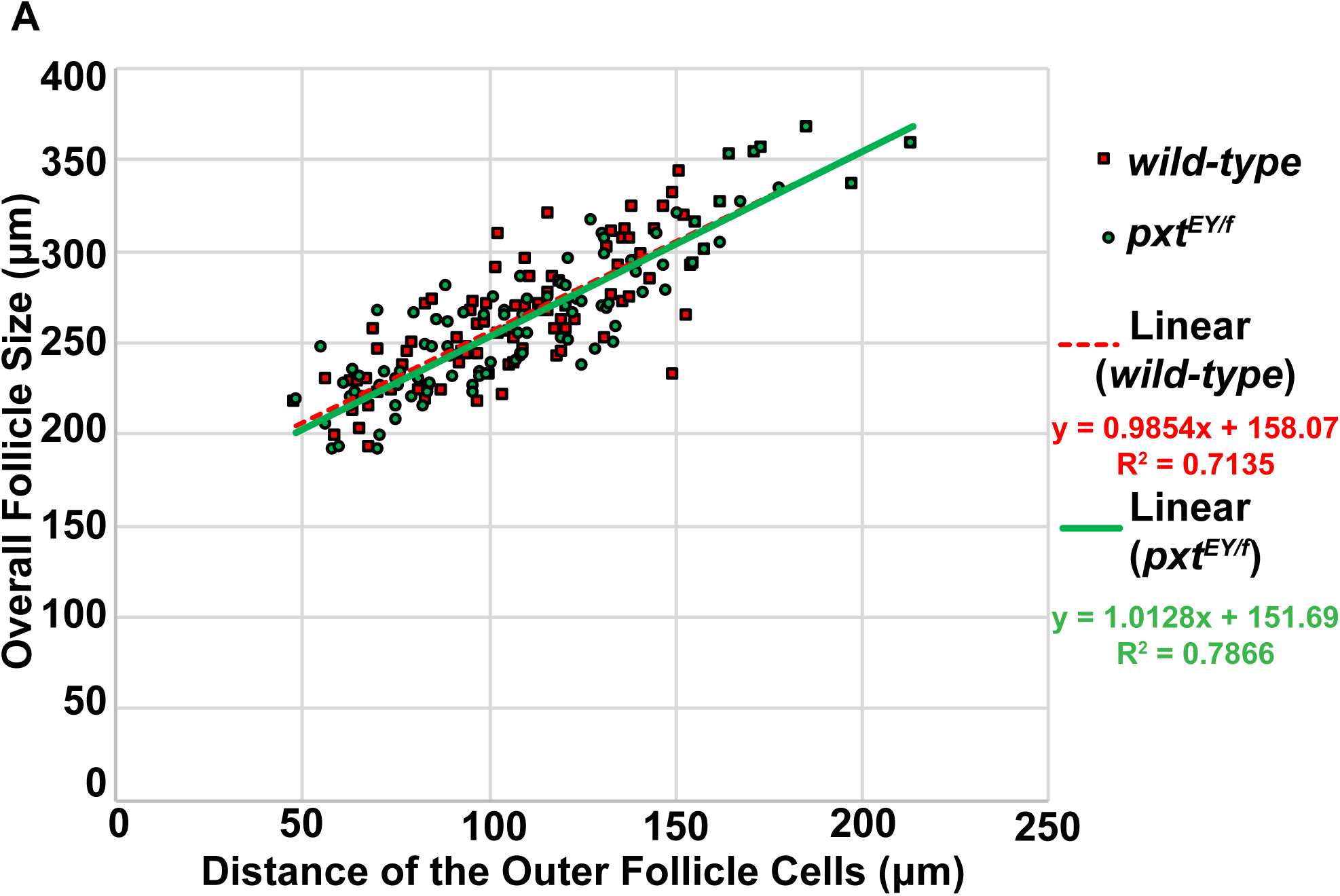
Outer follicle cell morphogenesis is normal when Pxt is lost. Graph of the follicle length vs outer follicle cell distance for wild-type (red) and *pxt^EY/f^* (green) follicles; each circle represents a single follicle and the best-fit lines provided. As the rate of change is similar between wild-type and *pxt* mutant follicles, this indicates that the outer follicle cell morphogenesis is normal in *pxt* mutants and therefore, the migration index can be used to assess border cell migration defects during S9.

**Supplementary Figure 4:**
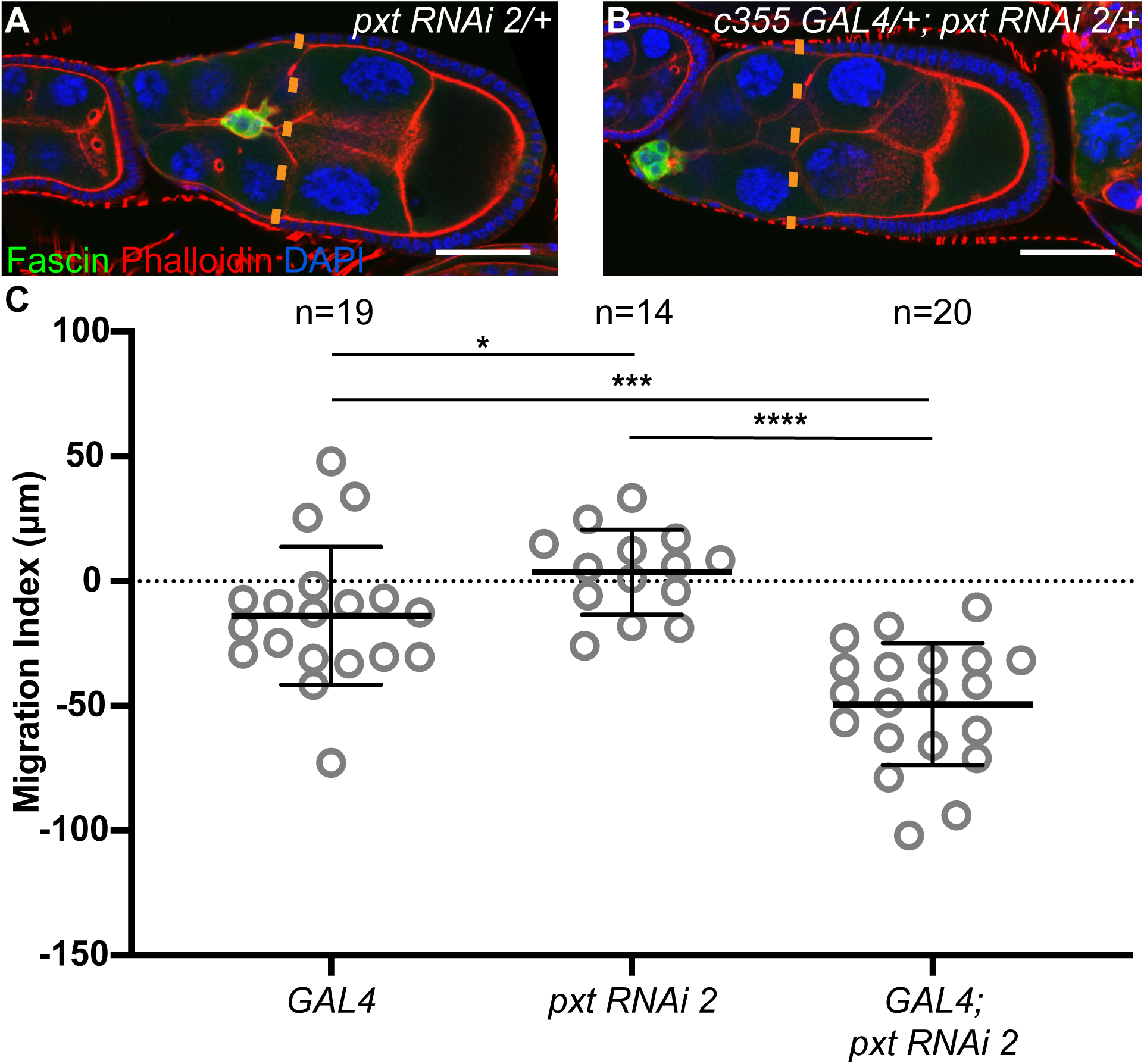
Somatic knockdown with a second *pxt* RNAi line results in delayed border cell migration during S9. A-B. Maximum projection of 3 confocal slices of S9 follicles of the indicated genotypes; anterior is to the left. A. Somatic GAL4 control (*c355 GAL4/+*). B. Somatic knockdown of *pxt* (c355 GAL4/+; *pxt RNAi 2/+*). Merged images: Fascin, green; Phalloidin (F-actin), red; and DAPI (DNA), blue. Orange dashed lines represent the position of the outer follicle cells. C. Graph of the migration index quantification during S9 for the indicated genotypes for two independent experiments. Each circle represents a single border cell cluster; the line indicates the average and the whiskers indicate the SD. Somatic knockdown of Pxt with a second RNAi construct (Vienna 104446) results in delayed border cell migration. ****p<0.0001, ***p<0.001, and *p<0.05. Scale bars= 50μm.

**Supplementary Figure 5:**
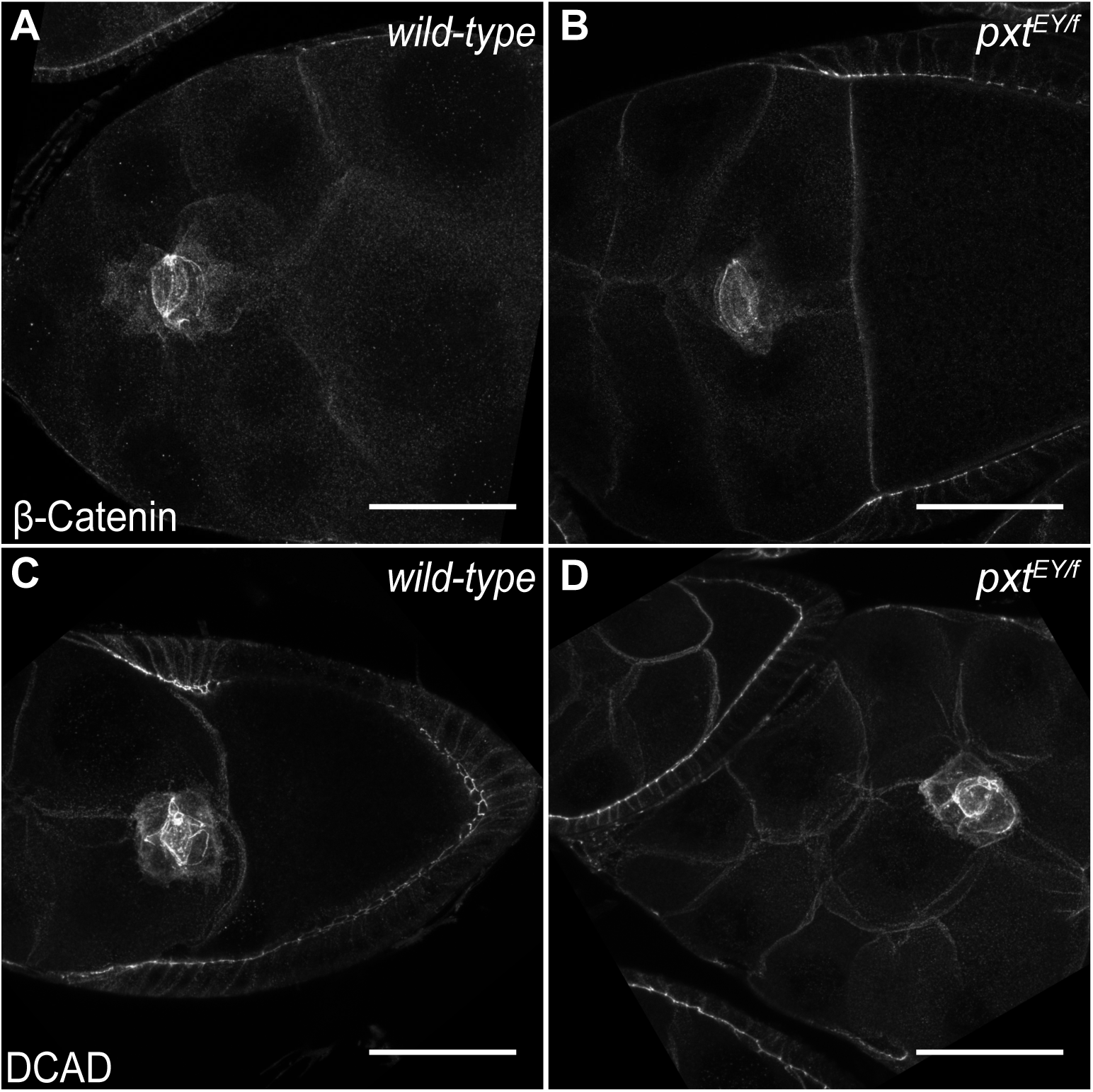
Pxt does not regulate β-Catenin and E-cadherin localization. A-D. Maximum projection of 3 confocal slices of S9 follicles of the indicated genotypes stained for β-Catenin (A-B) or *Drosophila* E-cadherin, DCAD (C-D); anterior is to the left. A, C. wild-type (*yw*). B, D. *pxt^EY/f^*. In both wild-type and *pxt* mutant follicles, β-Catenin and E-cadherin primarily localize to the border cell-border cell contacts. Scale bars= 50μm.

**Supplementary Figure 6:**
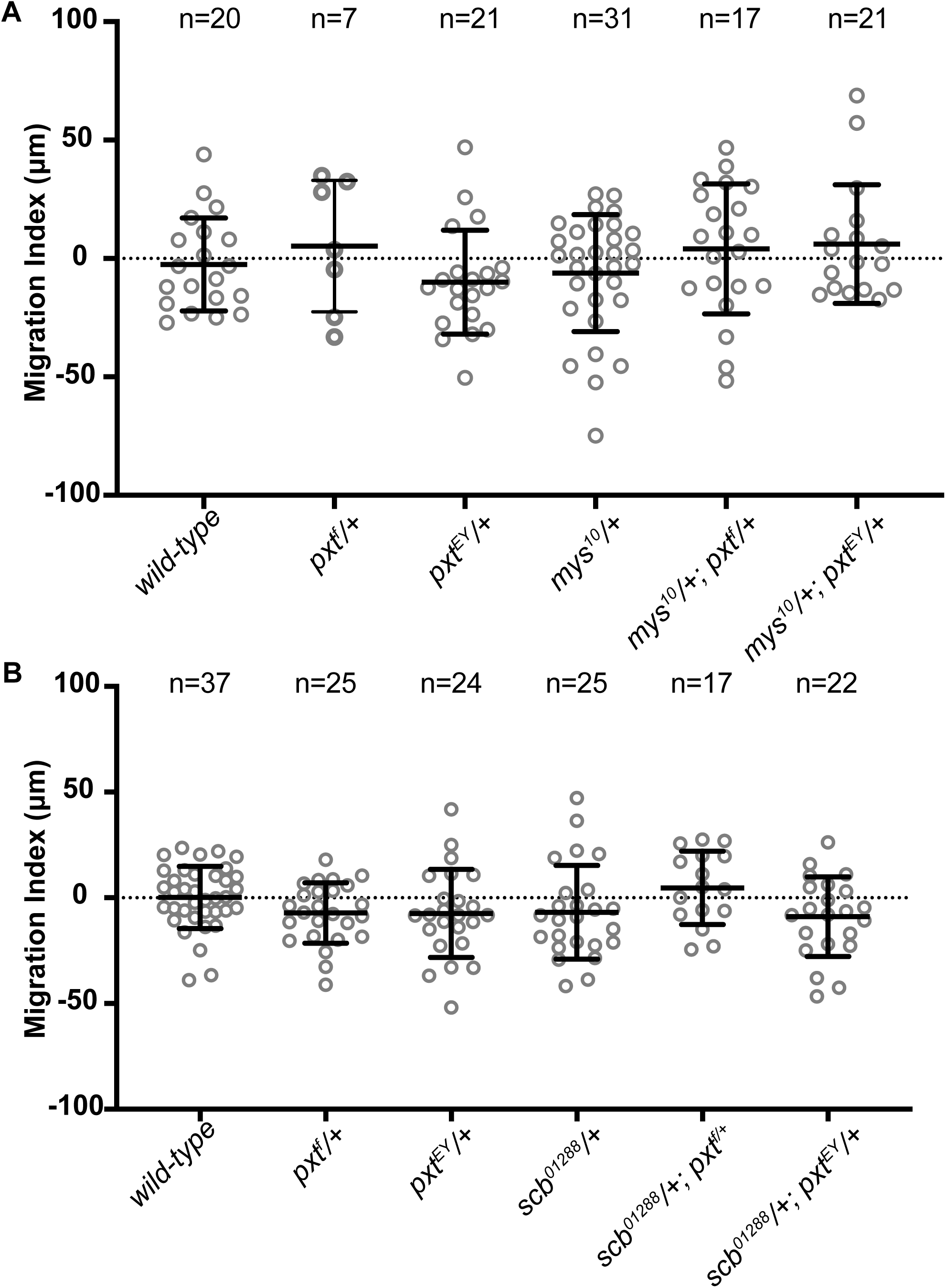
Dominant genetic interactions between *pxt* and integrin subunits. A-B. Graphs of the migration index quantification during Stage 9 for the indicated genotypes. Each circle represents a single border cell cluster; the line indicates the average and the whiskers indicate the SD. Heterozygosity for mutations in *β*PS-integrin (*mys^10^/+*), *α*PS3-integrin (*scb^01288^/+*), or *pxt* (*pxt^f^/+* or *pxt^EY^/+*), and double heterozygotes for *pxt* and an integrin subunit (*mys^10^/+; pxt/+* and *scb^01288^/+; pxt/+*) exhibit normal border cell migration.

## References

Abramoff, M.D., Magalhaes, P.J., and Ram, S.J. (2004). Image Proccesing with ImageJ. Biophotonics International 11, 36–42.

Alexander, S., and Friedl, P. (2012). Cancer invasion and resistance: interconnected processes of disease progression and therapy failure. Trends Mol Med 18, 13–26.

Anilkumar, N., Parsons, M., Monk, R., Ng, T., and Adams, J.C. (2003). Interaction of fascin and protein kinase Calpha: a novel intersection in cell adhesion and motility. EMBO J 22, 5390–5402.

Aranjuez, G., Burtscher, A., Sawant, K., Majumder, P., and McDonald, J.A. (2016). Dynamic myosin activation promotes collective morphology and migration by locally balancing oppositional forces from surrounding tissue. Mol Biol Cell 27, 1898–1910.

Bai, X., Wang, J., Zhang, L., Ma, J., Zhang, H., Xia, S., Zhang, M., Ma, X., Guo, Y., Rong, R., Cheng, S., Shu, W., Wang, Y., and Leng, J. (2013). Prostaglandin E(2) receptor EP1-mediated phosphorylation of focal adhesion kinase enhances cell adhesion and migration in hepatocellular carcinoma cells. Int J Oncol 42, 1833–1841.

Bai, X.M., Zhang, W., Liu, N.B., Jiang, H., Lou, K.X., Peng, T., Ma, J., Zhang, L., Zhang, H., and Leng, J. (2009). Focal adhesion kinase: important to prostaglandin E2-mediated adhesion, migration and invasion in hepatocellular carcinoma cells. Oncol Rep 21, 129–136.

Bianco, A., Poukkula, M., Cliffe, A., Mathieu, J., Luque, C.M., Fulga, T.A., and Rorth, P. (2007). Two distinct modes of guidance signalling during collective migration of border cells. Nature 448, 362–365.

Boyle, M., Bonini, N., and DiNardo, S. (1997). Expression and function of clift in the development of somatic gonadal precursors within the Drosophila mesoderm. Development 124, 971–982.

Brower, D.L., Wilcox, M., Piovant, M., Smith, R.J., and Reger, L.A. (1984). Related cell-surface antigens expressed with positional specificity in Drosophila imaginal discs. Proc Natl Acad Sci U S A 81, 7485–7489.

Cai, D., Chen, S.C., Prasad, M., He, L., Wang, X., Choesmel-Cadamuro, V., Sawyer, J.K., Danuser, G., and Montell, D.J. (2014). Mechanical feedback through E-cadherin promotes direction sensing during collective cell migration. Cell 157, 1146–1159.

Cant, K., Knowles, B.A., Mooseker, M.S., and Cooley, L. (1994). Drosophila singed, a fascin homolog, is required for actin bundle formation during oogenesis and bristle extension. J Cell Biol 125, 369–380.

Cha, Y.I., and DuBois, R.N. (2007). NSAIDs and cancer prevention: targets downstream of COX-2. Annu Rev Med 58, 239–252.

Cha, Y.I., Kim, S.H., Sepich, D., Buchanan, F.G., Solnica-Krezel, L., and DuBois, R.N. (2006). Cyclooxygenase-1-derived PGE2 promotes cell motility via the G-protein-coupled EP4 receptor during vertebrate gastrulation. Genes Dev 20, 77–86.

Cha, Y.I., Kim, S.H., Solnica-Krezel, L., and Dubois, R.N. (2005). Cyclooxygenase-1 signaling is required for vascular tube formation during development. Dev Biol 282, 274–283.

Chan, C., Jankova, L., Fung, C.L., Clarke, C., Robertson, G., Chapuis, P.H., Bokey, L., Lin, B.P., Dent, O.F., and Clarke, S. (2010). Fascin expression predicts survival after potentially curative resection of node-positive colon cancer. Am J Surg Pathol 34, 656–666.

Chen, W.S., Wei, S.J., Liu, J.M., Hsiao, M., Kou-Lin, J., and Yang, W.K. (2001). Tumor invasiveness and liver metastasis of colon cancer cells correlated with cyclooxygenase-2 (COX-2) expression and inhibited by a COX-2-selective inhibitor, etodolac. Int J Cancer 91, 894–899.

De Graeve, F.M., Van de Bor, V., Ghiglione, C., Cerezo, D., Jouandin, P., Ueda, R., Shashidhara, L.S., and Noselli, S. (2012). Drosophila apc regulates delamination of invasive epithelial clusters. Dev Biol 368, 76–85.

De Pascalis, C., and Etienne-Manneville, S. (2017). Single and collective cell migration: the mechanics of adhesions. Mol Biol Cell 28, 1833–1846.

Denkert, C., Winzer, K.J., Muller, B.M., Weichert, W., Pest, S., Kobel, M., Kristiansen, G., Reles, A., Siegert, A., Guski, H., and Hauptmann, S. (2003). Elevated expression of cyclooxygenase-2 is a negative prognostic factor for disease free survival and overall survival in patients with breast carcinoma. Cancer 97, 2978–2987.

Digiacomo, G., Ziche, M., Dello Sbarba, P., Donnini, S., and Rovida, E. (2015). Prostaglandin E2 transactivates the colony-stimulating factor-1 receptor and synergizes with colony-stimulating factor-1 in the induction of macrophage migration via the mitogen-activated protein kinase ERK1/2. FASEB J 29, 2545–2554.

Dinkins, M.B., Fratto, V.M., and Lemosy, E.K. (2008). Integrin alpha chains exhibit distinct temporal and spatial localization patterns in epithelial cells of the Drosophila ovary. Dev Dyn 237, 3927–3939.

Dormond, O., Bezzi, M., Mariotti, A., and Ruegg, C. (2002). Prostaglandin E2 promotes integrin alpha Vbeta 3-dependent endothelial cell adhesion, rac-activation, and spreading through cAMP/PKA-dependent signaling. J Biol Chem 277, 45838–45846.

Dormond, O., Foletti, A., Paroz, C., and Ruegg, C. (2001). NSAIDs inhibit alpha V beta 3 integrin-mediated and Cdc42/Rac-dependent endothelial-cell spreading, migration and angiogenesis. Nat Med 7, 1041–1047.

Fischer, J.A., Giniger, E., Maniatis, T., and Ptashne, M. (1988). GAL4 activates transcription in Drosophila. Nature 332, 853–856.

Friedl, P., and Gilmour, D. (2009). Collective cell migration in morphogenesis, regeneration and cancer. Nat Rev Mol Cell Biol 10, 445–457.

Friedl, P., and Mayor, R. (2017). Tuning Collective Cell Migration by Cell-Cell Junction Regulation. Cold Spring Harb Perspect Biol 9.

Gallo, O., Masini, E., Bianchi, B., Bruschini, L., Paglierani, M., and Franchi, A. (2002). Prognostic significance of cyclooxygenase-2 pathway and angiogenesis in head and neck squamous cell carcinoma. Hum Pathol 33, 708–714.

Giampieri, S., Manning, C., Hooper, S., Jones, L., Hill, C.S., and Sahai, E. (2009). Localized and reversible TGFbeta signalling switches breast cancer cells from cohesive to single cell motility. Nat Cell Biol 11, 1287–1296.

Grashoff, C., Hoffman, B.D., Brenner, M.D., Zhou, R., Parsons, M., Yang, M.T., McLean, M.A., Sligar, S.G., Chen, C.S., Ha, T., and Schwartz, M.A. (2010). Measuring mechanical tension across vinculin reveals regulation of focal adhesion dynamics. Nature 466, 263–266.

Groen, C.M., Spracklen, A.J., Fagan, T.N., and Tootle, T.L. (2012). Drosophila Fascin is a novel downstream target of prostaglandin signaling during actin remodeling. Mol Biol Cell 23, 4567–4578.

Harburger, D.S., and Calderwood, D.A. (2009). Integrin signalling at a glance. J Cell Sci 122, 159–163.

Hashimoto, Y., Shimada, Y., Kawamura, J., Yamasaki, S., and Imamura, M. (2004). The prognostic relevance of fascin expression in human gastric carcinoma. Oncology 67, 262–270.

Hoggatt, J., Singh, P., Sampath, J., and Pelus, L.M. (2009). Prostaglandin E2 enhances hematopoietic stem cell homing, survival, and proliferation. Blood 113, 5444–5455.

Jayo, A., Malboubi, M., Antoku, S., Chang, W., Ortiz-Zapater, E., Groen, C., Pfisterer, K., Tootle, T., Charras, G., Gundersen, G.G., and Parsons, M. (2016). Fascin Regulates Nuclear Movement and Deformation in Migrating Cells. Dev Cell 38, 371–383.

Khalil, A.A., Ilina, O., Gritsenko, P.G., Bult, P., Span, P.N., and Friedl, P. (2017). Collective invasion in ductal and lobular breast cancer associates with distant metastasis. Clin Exp Metastasis 34, 421–429.

Khuri, F.R., Wu, H., Lee, J.J., Kemp, B.L., Lotan, R., Lippman, S.M., Feng, L., Hong, W.K., and Xu, X.C. (2001). Cyclooxygenase-2 overexpression is a marker of poor prognosis in stage I non-small cell lung cancer. Clin Cancer Res 7, 861–867.

Kobayashi, K., Omori, K., and Murata, T. (2018). Role of prostaglandins in tumor microenvironment. Cancer Metastasis Rev 37, 347–354.

Lamb, M.C., Anliker, K.K., and Tootle, T.L. (2019). Fascin regulates protrusions and delamination to mediate invasive, collective cell migration *in vivo*. bioRxiv, 734475.

Li, H.J., Reinhardt, F., Herschman, H.R., and Weinberg, R.A. (2012). Cancer-stimulated mesenchymal stem cells create a carcinoma stem cell niche via prostaglandin E2 signaling. Cancer Discov 2, 840–855.

Li, X., Zheng, H., Hara, T., Takahashi, H., Masuda, S., Wang, Z., Yang, X., Guan, Y., and Takano, Y. (2008). Aberrant expression of cortactin and fascin are effective markers for pathogenesis, invasion, metastasis and prognosis of gastric carcinomas. Int J Oncol 33, 69–79.

Li, Y.J., Kanaji, N., Wang, X.Q., Sato, T., Nakanishi, M., Kim, M., Michalski, J., Nelson, A.J., Farid, M., Basma, H., Patil, A., Toews, M.L., Liu, X., and Rennard, S.I. (2015). Prostaglandin E2 switches from a stimulator to an inhibitor of cell migration after epithelial-to-mesenchymal transition. Prostaglandins Other Lipid Mediat 116–117, 1-9.

Liu, F., Mih, J.D., Shea, B.S., Kho, A.T., Sharif, A.S., Tager, A.M., and Tschumperlin, D.J. (2010). Feedback amplification of fibrosis through matrix stiffening and COX-2 suppression. J Cell Biol 190, 693–706.

Llense, F., and Martin-Blanco, E. (2008). JNK signaling controls border cell cluster integrity and collective cell migration. Curr Biol 18, 538–544.

Lyons, T.R., O’Brien, J., Borges, V.F., Conklin, M.W., Keely, P.J., Eliceiri, K.W., Marusyk, A., Tan, A.C., and Schedin, P. (2011). Postpartum mammary gland involution drives progression of ductal carcinoma in situ through collagen and COX-2. Nat Med 17, 1109–1115.

Mayor, R., and Etienne-Manneville, S. (2016). The front and rear of collective cell migration. Nat Rev Mol Cell Biol 17, 97–109.

Mayoral, R., Fernandez-Martinez, A., Bosca, L., and Martin-Sanz, P. (2005). Prostaglandin E2 promotes migration and adhesion in hepatocellular carcinoma cells. Carcinogenesis 26, 753–761.

Medioni, C., and Noselli, S. (2005). Dynamics of the basement membrane in invasive epithelial clusters in Drosophila. Development 132, 3069–3077.

Menter, D.G., and Dubois, R.N. (2012). Prostaglandins in cancer cell adhesion, migration, and invasion. Int J Cell Biol 2012, 723419.

Montell, D.J. (2003). Border-cell migration: the race is on. Nat Rev Mol Cell Biol 4, 13–24.

Montell, D.J., Rorth, P., and Spradling, A.C. (1992). slow border cells, a locus required for a developmentally regulated cell migration during oogenesis, encodes Drosophila C/EBP. Cell 71, 51–62.

Montell, D.J., Yoon, W.H., and Starz-Gaiano, M. (2012). Group choreography: mechanisms orchestrating the collective movement of border cells. Nat Rev Mol Cell Biol 13, 631–645.

Niewiadomska, P., Godt, D., and Tepass, U. (1999). DE-Cadherin is required for intercellular motility during Drosophila oogenesis. J Cell Biol 144, 533–547.

North, T.E., Goessling, W., Walkley, C.R., Lengerke, C., Kopani, K.R., Lord, A.M., Weber, G.J., Bowman, T.V., Jang, I.H., Grosser, T., Fitzgerald, G.A., Daley, G.Q., Orkin, S.H., and Zon, L.I. (2007). Prostaglandin E2 regulates vertebrate haematopoietic stem cell homeostasis. Nature 447, 1007–1011.

Oda, H., Uemura, T., Harada, Y., Iwai, Y., and Takeichi, M. (1994). A Drosophila homolog of cadherin associated with armadillo and essential for embryonic cell-cell adhesion. Dev Biol 165, 716–726.

Okada, K., Shimura, T., Asakawa, K., Hashimoto, S., Mochida, Y., Suehiro, T., and Kuwano, H. (2007). Fascin expression is correlated with tumor progression of extrahepatic bile duct cancer. Hepatogastroenterology 54, 17–21.

Pandya, P., Orgaz, J.L., and Sanz-Moreno, V. (2017). Modes of invasion during tumour dissemination. Mol Oncol 11, 5–27.

Patel, N.H., Snow, P.M., and Goodman, C.S. (1987). Characterization and cloning of fasciclin III: a glycoprotein expressed on a subset of neurons and axon pathways in Drosophila. Cell 48, 975–988.

Platt, J., and Michael, A. (1983). Retardation of fading and enhancement of intensity of immunofluorescence by p-phenylenediamine. J Histochem Cytochem 6, 840–842.

Prasad, M., and Montell, D.J. (2007). Cellular and molecular mechanisms of border cell migration analyzed using time-lapse live-cell imaging. Dev Cell 12, 997–1005.

Riggleman, B., Schedl, P., and Wieschaus, E. (1990). Spatial expression of the Drosophila segment polarity gene armadillo is posttranscriptionally regulated by wingless. Cell 63, 549–560.

Rolland, P.H., Martin, P.M., Jacquemier, J., Rolland, A.M., and Toga, M. (1980). Prostaglandin in human breast cancer: Evidence suggesting that an elevated prostaglandin production is a marker of high metastatic potential for neoplastic cells. J Natl Cancer Inst 64, 1061–1070.

Sales, K.J., Boddy, S.C., and Jabbour, H.N. (2008). F-prostanoid receptor alters adhesion, morphology and migration of endometrial adenocarcinoma cells. Oncogene 27, 2466–2477.

Scarpa, E., and Mayor, R. (2016). Collective cell migration in development. J Cell Biol 212, 143–155.

Speirs, C.K., Jernigan, K.K., Kim, S.H., Cha, Y.I., Lin, F., Sepich, D.S., DuBois, R.N., Lee, E., and Solnica-Krezel, L. (2010). Prostaglandin Gbetagamma signaling stimulates gastrulation movements by limiting cell adhesion through Snai1a stabilization. Development 137, 1327–1337.

Spracklen, A.J., Kelpsch, D.J., Chen, X., Spracklen, C.N., and Tootle, T.L. (2014). Prostaglandins temporally regulate cytoplasmic actin bundle formation during Drosophila oogenesis. Mol Biol Cell 25, 397–411.

Spracklen, A.J., Lamb, M.C., Groen, C.M., and Tootle, T.L. (2019). Pharmaco-Genetic Screen To Uncover Actin Regulators Targeted by Prostaglandins During Drosophila Oogenesis. G3 (Bethesda).

Spracklen, A.J., and Tootle, T.L. (2015). Drosophila – a model for studying prostaglandin signaling. In: Bioactive Lipid Mediators: Current Reviews and Protocols., eds. T. Yokomizo and M. Murakami: Springer Protocols, 181–197.

Spradling, A. (1993). Developmental genetics of oogenesis. In: The development of Drosophila melanogaster: Cold Spring Harbor Laboratory Press, 1–70.

Stuelten, C.H., Parent, C.A., and Montell, D.J. (2018). Cell motility in cancer invasion and metastasis: insights from simple model organisms. Nat Rev Cancer 18, 296–312.

Tootle, T.L. (2013). Genetic insights into the in vivo functions of prostaglandin signaling. Int J Biochem Cell Biol.

Tootle, T.L., and Spradling, A.C. (2008). Drosophila Pxt: a cyclooxygenase-like facilitator of follicle maturation. Development 135, 839–847.

Tootle, T.L., Williams, D., Hubb, A., Frederick, R., and Spradling, A. (2011). Drosophila eggshell production: identification of new genes and coordination by Pxt. PLoS One 6, e19943.

Tsujii, M., Kawano, S., and DuBois, R.N. (1997). Cyclooxygenase-2 expression in human colon cancer cells increases metastatic potential. Proc Natl Acad Sci U S A 94, 3336–3340.

Ugwuagbo, K.C., Maiti, S., Omar, A., Hunter, S., Nault, B., Northam, C., and Majumder, M. (2019). Prostaglandin E2 promotes embryonic vascular development and maturation in zebrafish. Biol Open 8.

Vicente-Manzanares, M., Choi, C.K., and Horwitz, A.R. (2009). Integrins in cell migration--the actin connection. J Cell Sci 122, 199–206.

Villari, G., Jayo, A., Zanet, J., Fitch, B., Serrels, B., Frame, M., Stramer, B.M., Goult, B.T., and Parsons, M. (2015). A direct interaction between fascin and microtubules contributes to adhesion dynamics and cell migration. J Cell Sci 128, 4601–4614.

Wang, D., and Dubois, R.N. (2010). Eicosanoids and cancer. Nat Rev Cancer 10, 181–193.

Wang, D., and DuBois, R.N. (2018). Role of prostanoids in gastrointestinal cancer. J Clin Invest 128, 2732–2742.

Yoder, B.J., Tso, E., Skacel, M., Pettay, J., Tarr, S., Budd, T., Tubbs, R.R., Adams, J.C., and Hicks, D.G. (2005). The expression of fascin, an actin-bundling motility protein, correlates with hormone receptor-negative breast cancer and a more aggressive clinical course. Clin Cancer Res 11, 186–192.

Zaccai, M., and Lipshitz, H.D. (1996). Differential distributions of two adducin-like protein isoforms in the Drosophila ovary and early embryo. Zygote 4, 159–166.

Zanet, J., Jayo, A., Plaza, S., Millard, T., Parsons, M., and Stramer, B. (2012). Fascin promotes filopodia formation independent of its role in actin bundling. J Cell Biol 197, 477–486.

